# Delineating organizational principles of the endogenous L-A virus by cryo-EM and computational analysis of native cell extracts

**DOI:** 10.1101/2022.07.15.498668

**Authors:** Lisa Schmidt, Christian Tüting, Fotis L. Kyrilis, Farzad Hamdi, Dmitry A. Semchonok, Gerd Hause, Annette Meister, Christian Ihling, Pranav N. M. Shah, Milton T. Stubbs, Andrea Sinz, David I. Stuart, Panagiotis L. Kastritis

## Abstract

The high abundance of most viruses in infected host cells benefits their structural characterization; endogenous viruses are present in low copy numbers, however, and are therefore challenging to investigate. Here, we retrieve cell extracts enriched with an endogenous virus, the yeast L-A virus. The determined cryo-EM structure discloses capsid-stabilizing cation-π stacking and an interplay of non-covalent interactions from ten distinct capsomere interfaces. The capsid-embedded mRNA decapping active site trench is supported by a constricting movement of two opposite-facing loops. tRNA-loaded polysomes and other biomacromolecules, presumably mRNA, are found in virus proximity while stacked dsRNA bundles and the sub-stoichiometric polymerase localize underneath the capsid surface. Mature viruses participate in larger viral communities resembling their rare in-cell equivalents in terms of size, composition, and inter-virus distances. Our results collectively describe a 3D-architecture of a viral milieu, opening the door to cellextract-based high-resolution structural virology.

## Introduction

Viruses typically consist of genetic material enveloped by a protein or a proteolipid shell^1^, promoting the transport of their genome from cell to cell. To proliferate, a virus depends on the host cell. A typical viral life cycle is generally divided into 6 steps^2^: attachment, penetration, uncoating, gene expression and replication, viral assembly, and, finally, release of progeny viruses, where the whole process starts anew. Viruses form viral factories during infections by hijacking the host cells’ replication machinery or even completely reprograming the host cell^3^. While many viruses inevitably lead to the host cell’s damage or death^4,5^, detrimental effects on the host’s health^6^ of endogenous viruses and retroviruses can sometimes be offset by neutral or even beneficial outcomes for their host^7^, *e.g*., shaping innate immune responses^6^.

Structural analysis of viruses is critical for a deeper understanding of their epidemiology, ecology and evolution. Virus diagnostics (including their detection, evaluation and handling) of cells are traditionally performed with low-resolution electron microscopy^8,9^ while intact virus structures at high resolution are studied with X-ray diffraction (crystallography and fiber diffraction) and cryogenic electron microscopy (single-particle cryo-electron microscopy (cryo-EM) or cryo-electron tomography (cryo-ET)) methods. Current advances in structural biology, especially in cryogenic electron microscopy^10^, have allowed an unprecedented analysis of viral architecture across methods and scales (*in silico*^11^, *in vitro*^12^, *in situ*^13^, and *in cellulo*^14^). This is exemplified by the current research on the ongoing COVID-19 pandemic^15^, in which viral research has deeply characterized the structure-function and interactions of SARS-CoV-2^16^, its life cycle^17^, and its multiple variants^18^. Most structural studies focus on exogenous viruses, not only because of their critical importance for understanding the molecular mechanisms of infection but also due to their intrinsic high abundance in the host cell that results in lysis (*e.g*., for SARS-CoV-2, an overall yield of ~10^5^ to 10^6^ virions has been observed per infected cell^19^). On the other hand, endogenous viruses that can integrate into the genome or be present within the host cells in low copy numbers are more challenging to study structurally^20^, especially within cells. In addition, the heterologous expression, characterization and evaluation of virus proteins generally include an overexpression step in a laboratory-grown cell, increasing the possibility of critical structural adaptation changes as well as limited recovery of virus interactions.

*S. cerevisiae*, a genetically well-described system, harbors in its genome an endogenous virus known as L-A virus^21^, a member of the Totivirus family, and its discovery was the main starting point of research on yeast virology^22^. The L-A virus is stably maintained in yeast cells and propagated vertically during mitosis or horizontally through cytoplasmic mixing during mating^23^ and appears to be symptomless for yeasts^24^. However, coinfections with satellite viruses - the M1, M2, M28, or Mlus^25^ which encode a toxic protein product - lead to the formation of an L-A helper/killer virus system^26^. The two viruses synergize with their host yeast for a killer phenotype, preventing contamination of the strain with other strains that lack the virus pair^26^. Additionally, another Totivirus, the L-BC virus, is often associated with an L-A virus infection.^27^ The L-BC virus is closely related to the L-A virus, with similar genomic size and similar life cycle, including capsid formation and polymerase activity, but without showing helper activity for the satellite viruses^28^.

The L-A virus contains a single 4.6 kb dsRNA^21^, encoding for a capsid protein (Gag), and the viral RNA-dependent RNA polymerase fusion protein (Gag/Pol)^29^. Due to ribosomal frameshifting^30^, the predicted Gag:Gag/Pol ratio is produced in 60:1^31,32^. The life cycle of the L-A virus takes place in the cytoplasm of the yeast cell: First, the plus strand is synthesized and transported outside the capsid. Then in the cytoplasm, ribosomes translate the RNA, and Gag, as well as Gag/Pol fusion proteins, will be formed. Gag will form an icosahedral capsid of 60 asymmetric dimers that include two Gag/Pol fusion protein copies, bound to the dsRNA^33^. Lastly, the minus-strand synthesis takes place, and the process begins anew **(Fig. 1a,** adapted from^34^). dsRNA viruses differ from other groups of viruses in that, their genome is never released from the capsid so that the latter performs not only a protective role but is also part of the mRNA synthesis and modification machinery^35^. Low-resolution single-particle cryo-EM analysis^36^ and a crystallographic structure of the capsid at 3.4 Å^37^ communicated more than 20 years ago demonstrated similarities of the capsid protein to Reoviridae^38^. These structures not only demonstrated that L-A virus Gag violates the tenets of quasi-equivalence by having 120 Gag copies^36,37^, but localized the first decapping enzyme described in the literature^39^. Recent studies have shown that the decapping site in both the L-A virus and the L-BC virus is able not only to uncap endogenous mRNAs for degradation decoy generation^40^, but also performs a ‘cap-snatching’ function^41,42^.

**Figure 1.**
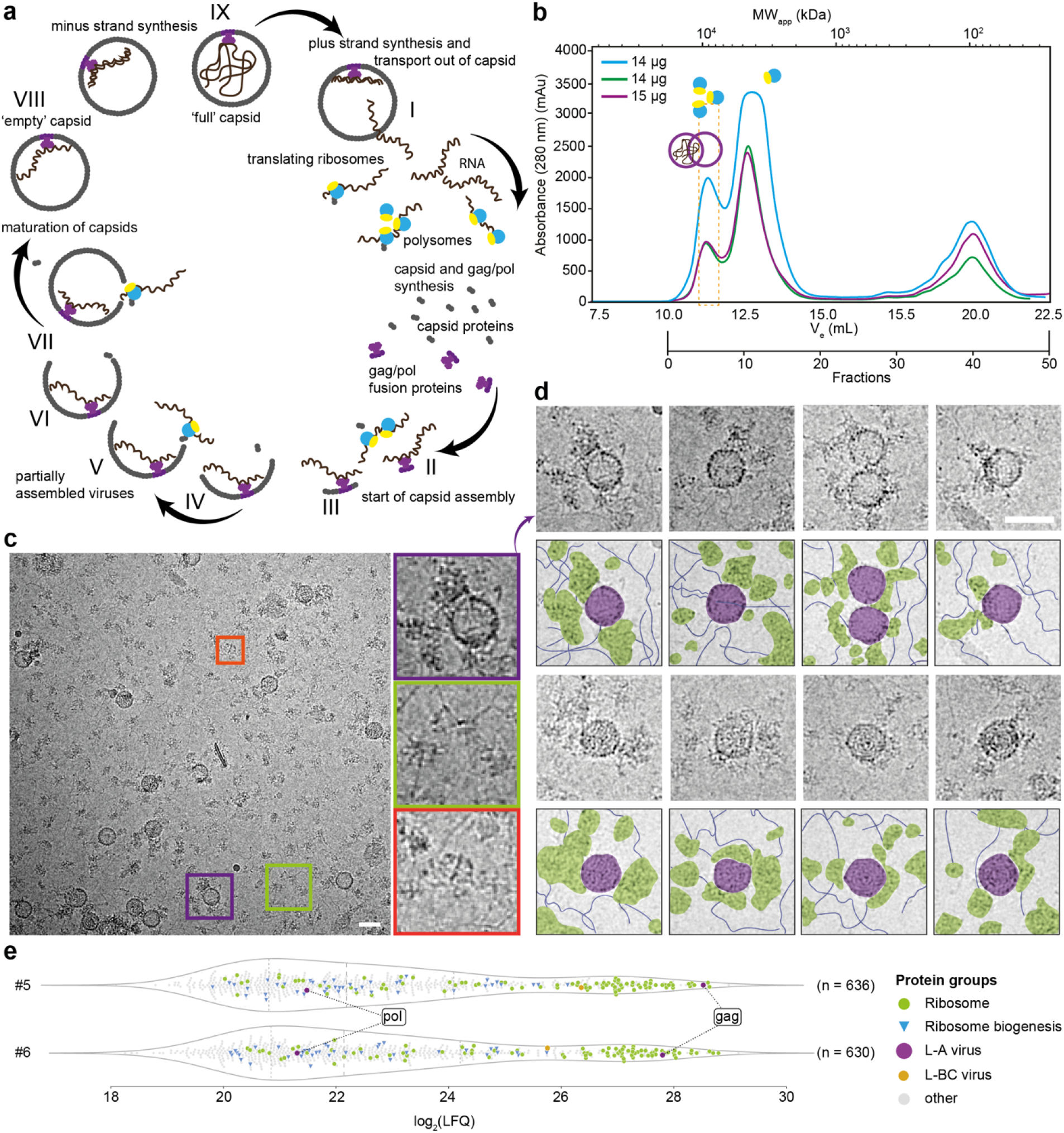
Identification of the L-A virus in native cell extracts: (a) Scheme of the yeast L-A virus life cycle (*states I-IX*). Replication starts with plus-strand synthesis and the transport out of the capsid (*state I*). Yeast ribosomes then translate the ssRNA and new viral particles (capsids) form (*states IIVII*). Last, the minus-strand is synthesized (*states VIII-IX*). Genome replication occurs in assembled viruses (*state IX*). (b) SEC profile of *S. cerevisiae* native cell extract. In this study, fractions 5 and 6, corresponding to ~10 MDa complexes, were further investigated (orange box). The purple circles show where full and empty L-A virus are expected based on relative retention. Accordingly, polysomes are also expected in the first peak and monosomes in the second peak. (c) Representative micrograph of fraction 5. A heterogeneous mixture of biomolecules can be seen, for example, the fatty acid synthase (red box), ribosomes (green box), and the L-A virus (purple box). (d) Representative crops of the L-A helper virus are shown in the vicinity of ribosomes, fibrillar structures, and other proteins. (e) LFQ-based MS quantification of fractions 5 and 6 to identify the LA helper virus and the fraction contents. Scale bars represent 50 nm.

In this work, we utilize the L-A helper virus as a model system for subsequent structural analysis after successful biochemical enrichment. The advantage of this approach is a closer-to-native observation of the endogenous virus because cell extracts are (a) less complex as compared to a whole-cell lysate or a cell but still retain principles of cellular organization^43,44,45^; (b) easily accessible to biochemical, biophysical, biocomputational and structural methods that can be combined with modern artificial-intelligence-based methods^46^; and (c) amenable to a selective increase in protein concentrations as a traditional biochemical specimen^43^. Our results collectively characterize the 3D ‘architecture’ of the endogenous L-A virus milieu, opening the door to structural virology in native cell extracts.

## Results

### The yeast L-A virus is identified in a cryo-EM-accessible eukaryotic cell extract

To isolate complexes of similar molecular weight within the native cell extract, we performed size exclusion chromatography (SEC) and reproducibly collected the resulting fractions in less than 8 hours from harvesting yeast cells (**Fig. 1b, Supplementary Fig. 1a, Methods**). The method, adapted from fractionation experiments on the thermophilic mold *Chaetomium thermophilum*^44^ with minor modifications involving changes in starting volume of cells and the process of resuspending the starting material before lysis (**Methods**), resulted in retrieval of high in-fraction protein concentrations (**Supplementary Fig. 1b**) amenable to direct cryo-EM analysis. After SEC, we vitrified fractions 5 and 6 that include megadalton (MDa) assemblies (**Fig. 1b,** orange dotted line). In the acquired cryo-EM micrographs, large unperturbed complexes are visible, including fatty acid synthase (FAS)^47^ (**Fig. 1c**) and polysomes (**Fig. 1c**) as well as prominent spherical signatures (**Fig. 1c**). A frequent observation while analyzing the micrographs was the reoccurring proximity of the spherical signature with filamentous structures and potential polysomes **(Fig. 1d)**.

We collected 7011 movies at a pixel size of 1.57 Å and performed image analysis of the derived spherical signatures (**Supplementary Fig. 2**), resulting in an icosahedrally averaged cryo-EM map at 3.77 Å resolution (FSC=0.143, **Supplementary Table 1**) in which an initial backbone trace could be built. A search for structural relatives in the Protein Data Bank (PDB)^48^ utilizing distance-based matrix alignments^49^, identified the (only available) 3.5 Å *Saccharomyces cerevisiae* L-A virus crystal structure^37^ as a potential match. Mass spectrometry (MS)-based protein identification of the fractionated cell extract verified the abundance of the L-A helper virus (Gag, Gag/Pol), and also the presence of the L-BC virus, albeit with lower abundance, after adding these endogenous protein sequences to the available yeast proteome (UP000002311)^50^ (**Fig. 1e, Supplementary Fig. 3a**).

To annotate the proteomic content of the studied extract, an analysis of the measured biological triplicates of the two L-A virus-containing fractions was performed. Ribosomal proteins were identified with very high abundance after streamlining the classification of identified proteins using the Kyoto Encyclopedia of Genes and Genomes (KEGG) pathway database^51^ **(Fig. 1e, Supplementary Fig. 3a)**. The capsid (gag) and the polymerase (pol) of the L-A virus were also identified within these fractions with high abundance **(Fig. 1e, Supplementary Fig. 3a)**. Calculation of the capsid:pol ratio per fraction from the MS data as performed for the endogenous pyruvate oxidation metabolon^44^ show an average of 1.8 +/- 0.49 pol present per capsid, in agreement with previous predictions based on frameshifting efficiency of the gag open reading frame^31^. Compared to other studies, we succeeded in (a) enriching the L-A virus within rapidly retrieved yeast cell extracts directly amenable for cryo-EM; (b) probing the “environment” and components of the L-A virus by quantitative mass spectrometry; and (c) identifying unambiguously the L-A virus using the high-resolution features of the calculated cryo-EM map by Cα-trace model building.

### The high-resolution cryo-EM structure of the L-A virus from a cell extract unveils structural and functional adaptations of the native state

A high-quality *de novo* model was built in the derived 3.77 Å (FSC=0.143, **Supplementary Fig. 3b**) L-A virus cryo-EM map (**Supplementary Table 1, Fig. 2a**) after image processing **(Supplementary Fig. 2)**, explaining well the densities **(Supplementary Fig. 3c)** that exhibited main-chain, and, often, side-chain resolution **(Fig. 2b)**. The icosahedral protein shell has the triangulation number T=2 formed by 120 protomers, with overall diameter (400 Å) and thickness (46 Å) comparable to its crystallographic counterpart, as well as a similar opening diameter at the icosahedral fivefold axes (18 Å) that serves as a gate for the exit of viral mRNA and entry of nucleotide triphosphates. This highly conserved overall L-A virus architecture that is retained in both cell extracts and in the crystal points to high stiffness of the capsid and its openings, indicating critical structural rigidity.

**Figure 2.**
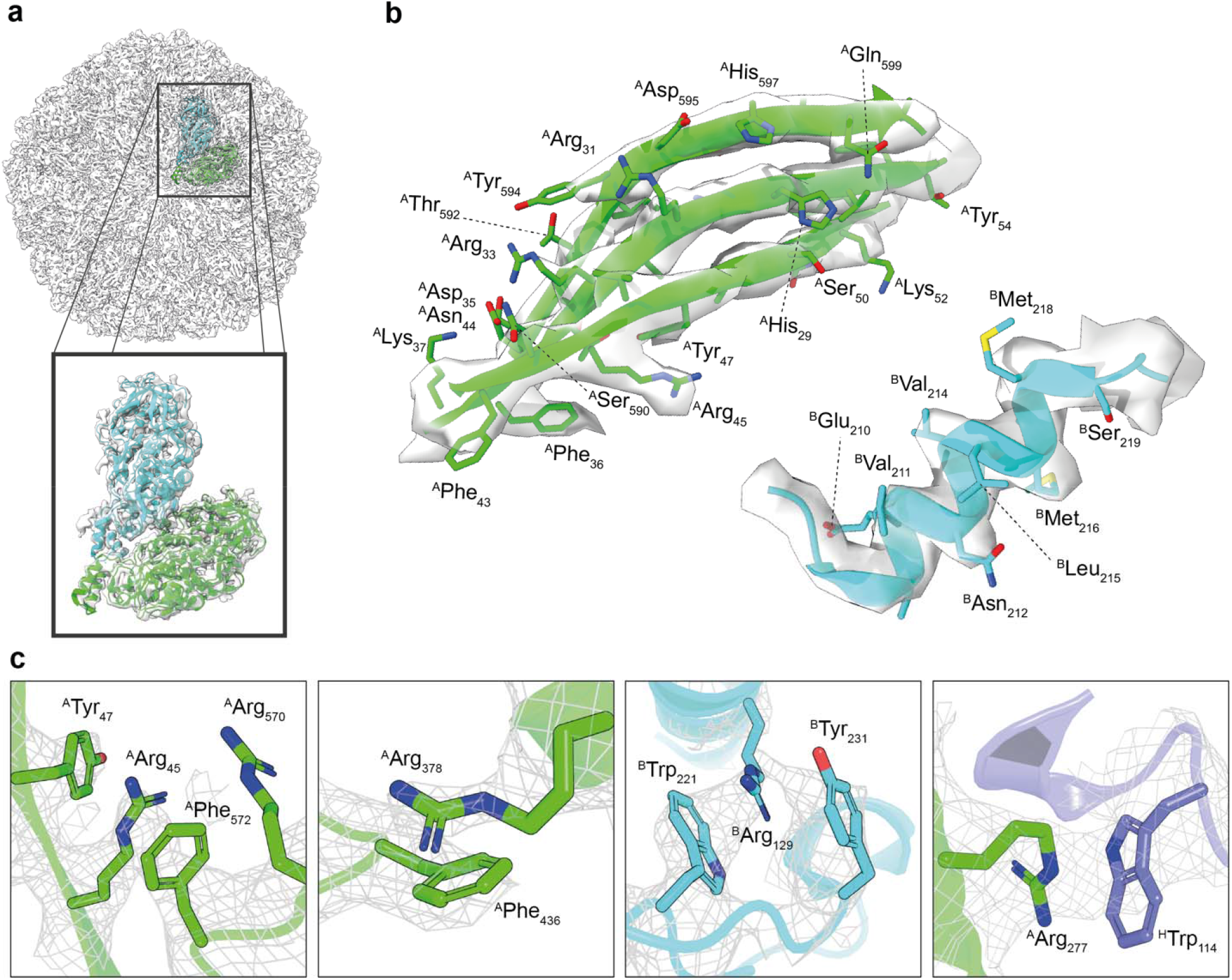
Distinctive features stabilizing the L-A capsid: (a) Derived cryo-EM map and model of the asymmetric subunit; Fitting is shown magnified in the insert. (b) Representative resolution features of the atomic model fitted in the cryo-EM map. In green, a close-up of a β-sheet is shown with the separation of the single strands clearly visible in the density of the map; in blue, an α-helix is shown with helical pitch visible at the resolution of 3.78 Å; side chain density is often reliable. (c) Representative cation-π interactions are shown between arginine and tyrosine, tryptophan or phenylalanine within chain A (green), chain B (blue), or between chain A and an adjacent chain B (slate).

Detailed analysis of the L-A virus identified previously undescribed stabilizing cation-π interactions in the capsid that mediate interactions within and also across LA protomers (**Fig. 2c**). In the asymmetric unit, 19 side chains are involved in forming 8 distinct cation-π interaction networks, formed within monomeric and dimeric interfaces (examples are shown in **Fig. 2c**). Remarkably, residues involved in these interactions are better resolved compared to side chains of the same type not involved in cation-π interactions (**Fig. 2c, Supplementary Fig. 3d, Supplementary Fig. 3e)**. The cation-π interaction represents a strong non-covalent bond^52^, with examples including (a) the structural presence of a single cation-π interaction in bacteriophage T5 that persists independently of capsid conformational change^53^ and (b) the prediction of another cation-π interaction in a dsRNA virus (bluetongue virus) after mutagenesis^54^. We deem the extended network of cation-π interactions observed to be highly specific (*e.g*., compared to non-specific hydrogen-bonding pattern) because a delocalized π-electron system must organize in the correct plane towards a cationic moiety. In the context of the L-A helper virus, cation-π interactions may contribute to both the folding of the monomeric building block and to the higher-order assembly of the virus itself.

The crystallographically determined regions of the capsomere exhibit substantial conformational differences between the two monomers in the resolved asymmetric unit^37^, *i.e*., residues N8–K12, G82, T96–I99, I111–T112, and G387–D396, which are also observed in the cryo-EM model. In addition, the latter reveals specific cell extract-based adaptations, distinct from those stemming from the dense crystal packing capsid conformation^37^ **(Supplementary Fig. 4a)**. Overall, the cryo-EM-derived asymmetric homodimer exhibits superior fits for both backbone (CC_cryo-EM_=0.85 vs. CC_x-ray_=0.79) and side-chain (CC_cryo-EM_=0.84 vs. CC_x-ray_=0.79) conformations **(Supplementary Fig. 4a)**. Structural changes compared to the crystal structure are detected in both capsomere subunits (**Supplementary Fig. 4b)**, specifically in regions E92–V104, G387–S393, and F525–T535, and regions A494–D505 of protomers A and B, respectively – all directly involved in capsid assembly (**Fig. 3a-d**). In detail, the protomer helix-turn-helix fold region E92–V104 forms the functional pores at the icosahedral five-fold axes (**Fig. 3b,** purple-blue), supported by the exact same region in protomer B **(Fig. 3b,** purple-blue). Such a dual role (supporting and pore-forming) for this minimal fold may rationalize the observed flexibility in the cryo-EM structure. Next, loop G387-S393 is buried in the dimeric interface of the capsomere (**Fig. 3c,** red) while loop F525–T535 is involved in higher-order binding of the that involve loop-loop interactions (**Fig. 3d,** blue). The flexible loop A494–D505 in capsomere protomer B interfaces with loop G387–S393, thereby further stabilizing the capsid (**Fig. 3d,** yellow).

**Figure 3.**
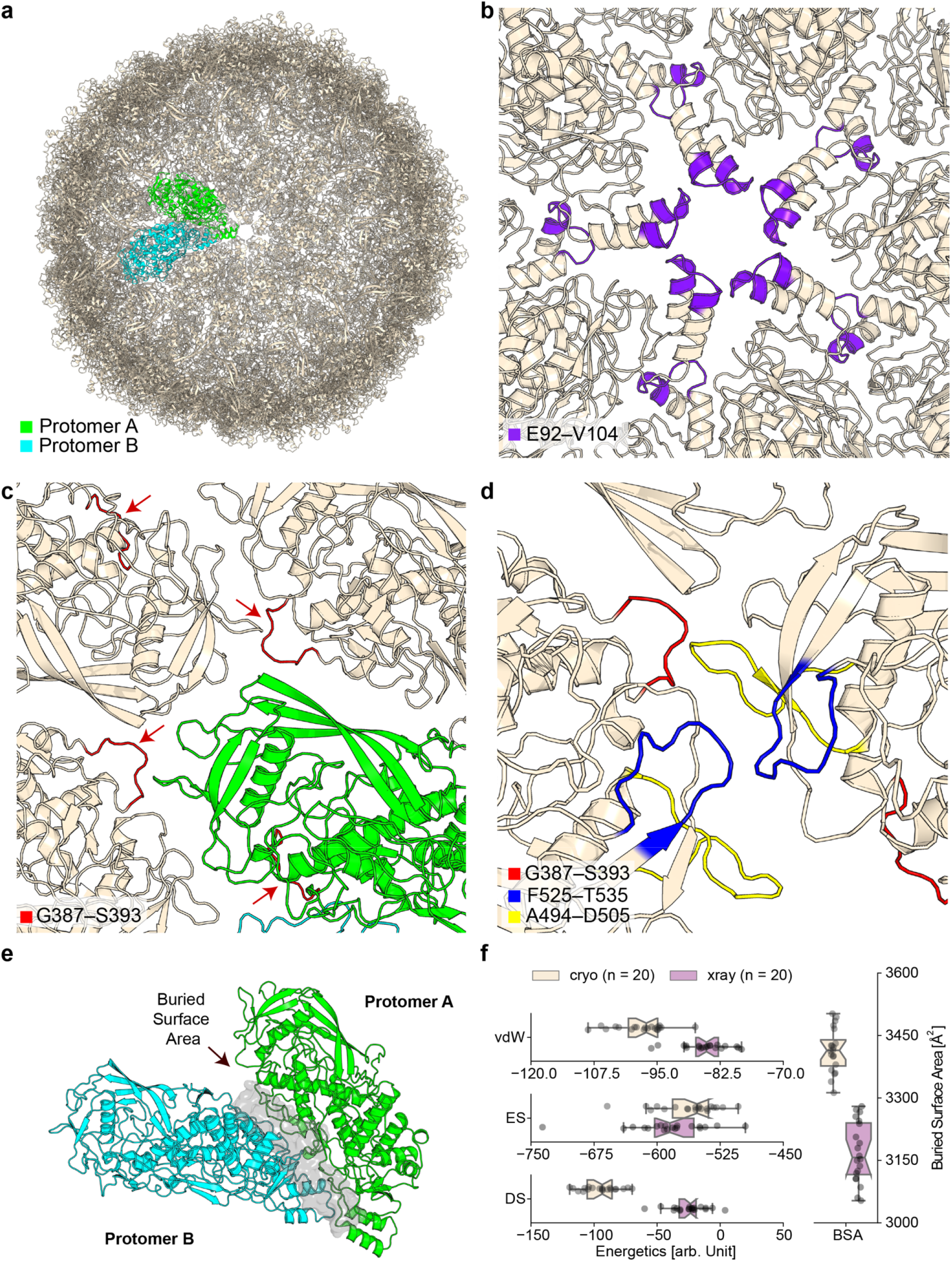
Structural plasticity of the L-A virus cryo-EM structure: (a) Overview of full capsid structure and the asymmetric unit (capsomere) composed of protomer A and B (b) protomer’s helix-turn-helix fold at region E92–V104 localizing at the five-fold axis, structuring and supporting the mRNA exit tunnel/nucleotide entry tunnel (c) loop G387–S393 is buried in the capsomeres dimeric interface and shown with red arrows (d) loop F525–T535 is involved in higher-order binding of the capsomeres communicating with other regions of identified flexibility, involving loops A494–D505 and G387–S393. (e) View of the capsomere composed of the two protomers A and B, highlighting the extensive surface area buried, forming the capsomere interface (3400 Å^2^). Chain A is colored in peach, chain B in mint green, and the interface in grey. (f) Scoring components derived for the interface after refinement of the capsomere shown in (e) calculated with the macromolecular modelling software HADDOCK. Non-covalent interactions calculated include van der Waals (VdW), electrostatics (ES), and desolvation energy scores (DS), all in arbitrary units (a.u.), and buried surface area (BSA), in Å^2^.

Overall, the above-mentioned distinct flexibility differences manifest in quantifiable energetic contributions calculated with the macromolecular modelling software HADDOCK^55^. The embedded capsomere interface **(Fig. 3e)** buries a large surface of 3400 Å^2^ (**Fig. 3f**) with electrostatics making major contributions (**Fig. 3f**). Extending energetic calculations to all proximal interfaces of the capsomere reveals interfaces of highly diverse energetic contributions where an interplay of van der Waals, electrostatics, and desolvation energies uniquely define each of the 9 surrounding interfaces as well (**Supplementary Fig. 5a**). Using these 9 unique interface energetic calculations, we propose a model of capsomere stability, which consists of 86 distinct steps of monomeric association to complete the capsid (**Supplementary Fig. 5b; Supplementary movie 1**).

### The mRNA decapping site trench shape is mediated by two flexible loops and is proximal to a potential protein-protein interaction site

mRNA decapping activity is widespread across dsRNA virus families but its location across protomers is not conserved^35^. In both the x-ray and cryo-EM L-A virus structures, region Gln139-Ser182 (containing the active site His154) (**Fig. 4a-d**), contributes to the outer capsid surface, mediates cellular mRNA decapping, and transfers the 7-methyl-GMP (m^7^ GMP) cap from the cellular mRNA 5’-end to the viral RNA 5’-end, countering a host exoribonuclease that targets uncapped RNAs. This site therefore directs competition for the use of the translation machinery. This active site was previously described to form a trench^26^ that involves residues located in loop regions. These loops retain similar conformations in the cryo-EM resolved structure, and, therefore, a similar local Coulomb surface potential is calculated (**Fig. 4a, 4b**). However, an additional distal pair of loops facing each other (regions P449-Y452 and T531-T535, the latter also involved in flexible region F525–T535 **(Fig. 3d)**) influences the diameter of the trench pore. Their distance is 3 Å closer in the cryo-EM structure, which is enough for H532 to close the trench (**Fig. 4c, d**). Although the presence of the open state cannot be dismissed in the endogenous state, this more occluded trench is found in the derived average, pointing to its prominence. Results disclose that a selectively open active site conformation is not common across all capsomeres, but, logically, must exist in proximity to the pore at the 5-fold axis; however, icosahedral averaging does not allow quantifying open and closed active site states.

**Figure 4.**
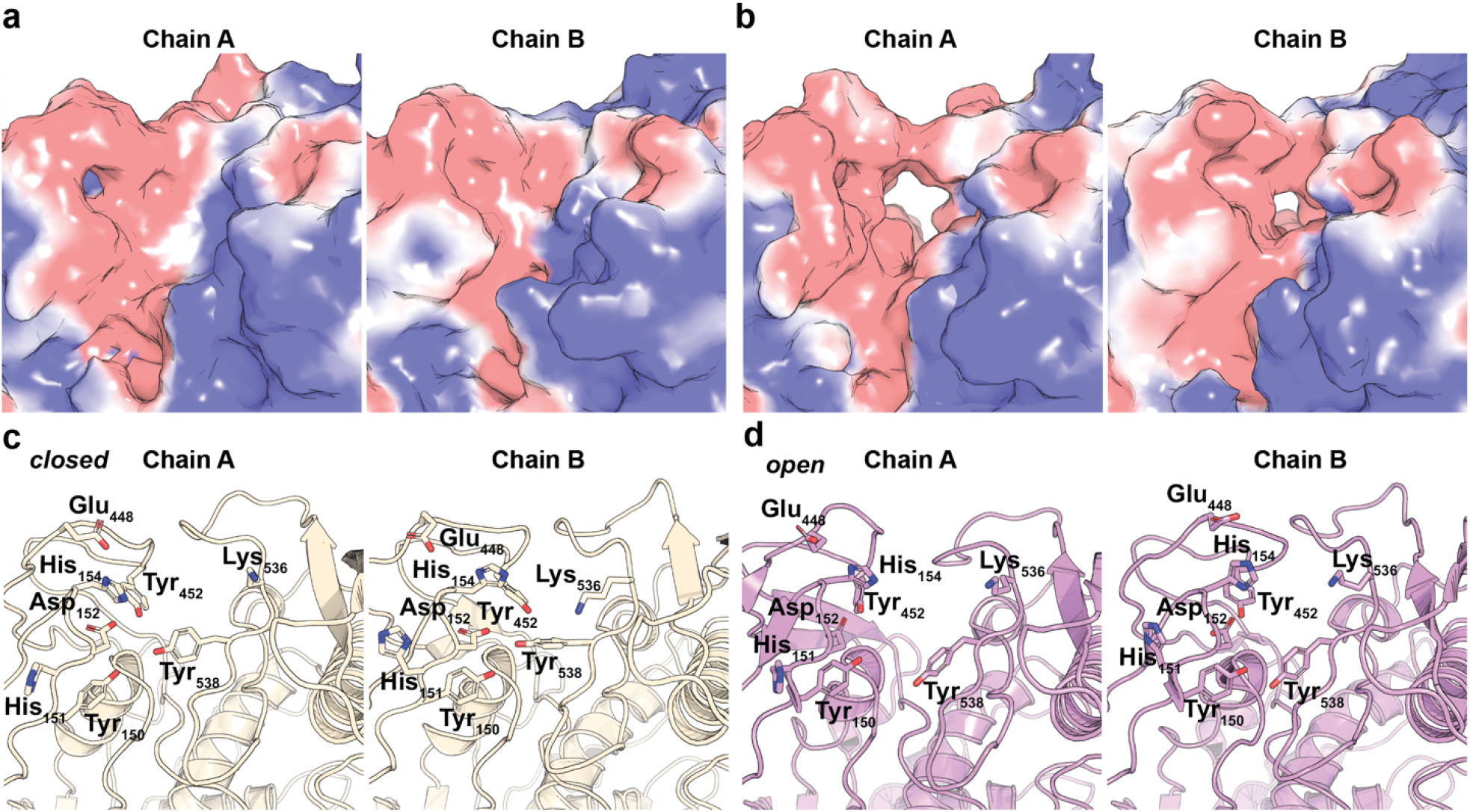
The mRNA decapping site trench shape is mediated by two flexible loops: (a-d) show the decapping site in surface (a, b) and cartoon (c, d) representations. Panels (a) and (c) show the conformations identified in the reported cryo-EM structure of the L-A virus capsid, whereas panels (b) and (d) illustrate those reported in the crystal structure. Chain A and Chain B correspond to the capsomeres subunits A and B respectively. Overall, the decapping center of the L-A virus capsid is identified in an occluded state, as seen in the cryo-EM data. The decapping center in the x-ray data is shown in an open state. Regions Gln139-Ser182 contribute to the outer capsid surface forming a trench assembled by flexible loops. His154 is found as the active residue for RNA decapping and is shown in (c) and (d).

Structural alignment of the structured region proximal to the active site (residues 491-587) using the DALI webserver^56^ identified domains with structural homology despite very low pairwise sequence identity (<10%) (**Supplementary Table 2, Supplementary Fig. 6a-b**). The structural domain comprises a minimal fold composed of a surface-exposed antiparallel β-sheet formed by 3 (2 long and 1 short) β-strands and inner α-helical and loop elements, buried in the protomer, and involves the two critical residues Tyr538 and Asp540 that participate in the mRNA active site decapping trench (**Fig. 4b, 4d**). Interestingly, this conserved minimal fold is analogous to the methyltransferase N6AMT and the cytotoxic nuclease domain of immunity protein 5 (im5)^57^ with a main-chain root-mean-square deviation (*rmsd*) reported by DALI of 3.2 and 2.8 Å, respectively (**Supplementary Table 2**). Overall, our results show an alternative conformation of the mRNA active site decapping trench as well as distant structural homologs with potential evolutionary implications.

### Asymmetric reconstruction of the L-A virus reveals tubular densities inside the capsid with implications for RNA-dependent RNA polymerase localization

To further characterize the L-A virus within cell extracts, we acquired 10067 movies from the fractionated extract at a lower pixel size (3.17 Å) to increase the number of imaged single particles. The lower-resolution icosahedrally-reconstructed cryo-EM map (**Supplementary Fig. 7a**) included 103214 particles in total with included 2D class averages that covered different views across the 2-fold, 3-fold, and 5-fold axes (**Supplementary Fig. 7b**), and reached Nyquist at 6.4 Å (FSC=0.143, **Supplementary Fig. 7c**) with high signal-to-noise ratio as shown by the map x,y slices (**Supplementary Fig. 7d).** After particle subtraction to eliminate the prominent capsid density, subsequent 2D classification showed mostly empty viruses (or viruses with averaged inner signal) as well as capsids with distinct inner densities (**Fig. 5a**). Utilizing 17549 density-subtracted particles that showed prominent inner density features, we derived an asymmetric reconstruction of the inner densities at ~18 Å (FSC=0.5, **Supplementary Fig. 7e, Fig. 5b**). The reconstructed inner density is asymmetrically positioned in the virus capsid (**Fig. 5b**), emanating below the 18 Å diameter opening at the 5-fold axis. The interior density clearly shows tubular, stacked densities resembling canonically packed dsRNA (**Fig. 5c**).

**Figure 5.**
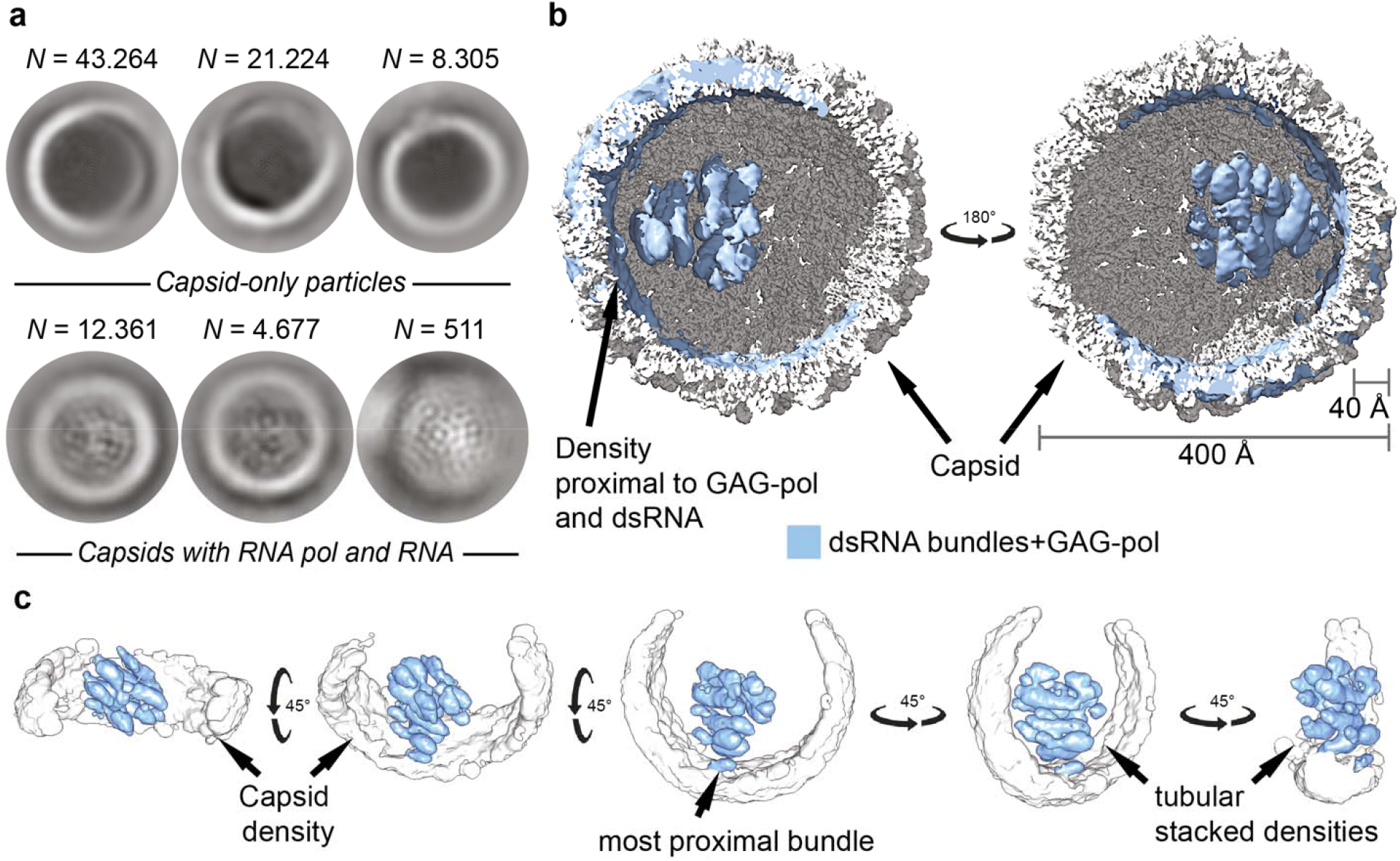
View inside the yeast L-A helper virus retrieved from native, endogenous yeast extracts: (a) 2D classification of inner densities of “full” L-A virus particles after subtracting the capsid density with CRYOSPARC. 2D class averages with capsid-only particles were discarded, and 2D class averages containing particles with internal density were used for 3D reconstruction. (b) 3D reconstruction of inner densities, colored light blue, superimposed on the capsid map (light grey). Arrowheads point at the capsid density and the most proximal inner density bundle. (c) Different views of the reconstructed inner density. Appearing capsid densities colored light grey and inner densities colored light blue shown in different views rotated 45° around the x and y axes.

The structure of the RNA-dependent RNA polymerase of the L-A virus is currently unknown and cannot be reliably placed at this resolution due to its rare presence and sub-stoichiometric abundance. Nevertheless, low-resolution densities proximal to the capsid were found (**Fig. 5c**), as well as densities inside the pentameric face pore (**Supplementary Fig. 8a)**. To delineate its relative positioning in the capsid interior, we employed artificial intelligence (AI)-based structural modelling^58^ of the 1505 amino acid gag-pol fusion product that is synthesized upon ribosomal frameshifting. Results show a high-quality atomic model for the individual gag and pol domains, within experimental error per each modelled fold (**Fig. 5d**). Specifically, an AI-based model of the pol domain was of high quality (**Supplementary Fig. 8b-c, 9a**) because plenty of experimentally determined structural templates currently exist in PDB^48^ (**Supplementary Fig. 9b).** Interestingly, ambiguously modelled regions include loops and specifically the positively-charged unstructured linker of the gag *C-ter* and the pol *N-ter* (**Supplementary Fig. 8b**).

Although this linker is 62 residues long, the Alphafold model places the pol domain interacting with the gag surface that would be exposed to the capsid interior (**Supplementary Fig. 8b**). Such interaction is indeed possible when looking at the complete L-A virus (**Supplementary Fig. 8b**). The linker region, which is highly charged could play a role in positioning the pol relative to the gag, possibly folding around the ordered pol molecule in a compacted state (**Supplementary Fig. 8c,** light pink color). Standard physical-chemical calculations for the disordered linker considering its calculated molecular weight and length suggest the following distances separating gag_C-ter_ and pol_N-ter_: In the unlikely fully extended linker conformation, the theoretical gagC-ter/polN-ter maximum distance is 235 Å (**Supplementary Fig. 8c**, darker pink color). However, in the more likely scenarios, *e.g*., if the linker is completely folded^59^, gag_C-ter_/pol_N-ter_ distance is ~30 Å, and in other possible structural states, such as molten globule, unfolded, and/or disordered^60^, gag_C-ter_/pol_N-ter_ distances correspond to ~35-45 Å. Consequently, results show that pol primarily explores proximal distances below the pore to the five-fold axis, coinciding with the proximity of multiple resolved tubular densities (**Fig. 5c**). These calculations imply that RNA replication within the L-A virus is confined to an asymmetric nano compartment with a < 50 Å potential size per gag-pol protomer, also supported by the linker’s complementary charge to dsRNA.

### Detecting higher-order interactions of the L-A virus and minimal communities

Analysis of the lower pixel size (3.17 Å) cryo-EM acquisition to improve statistics (10067 movies) shows the systematic proximity of electron-dense material to pleomorphic L-A viruses (**Supplementary Fig. 10a**) which we subsequently analyzed (**Supplementary Fig. 10b-f**). By 2D averaging picked L-A virus particles, 9711 single-particles with proximal structural signatures can be derived, although the direct proximity must be highly flexible given the diffused densities (**Supplementary Fig. 10b**). In addition, a 3D reconstruction of a ribosome, resolved at 8.1 Å (unmasked, FSC=0.143) from 627748 particles (FSC=0.143, **Supplementary Fig. 11a, Supplementary Table 1**) corroborates the high abundance of ribosomes in the same fractions with the L-A virus. Overall, the 2D classification of picked single particles unambiguously shows that the cell extract fraction contains polysomes (**Supplementary Fig. 11b**) with a 3D conformation resembling published polysome conformations^61^ and stalled disome states^62^ (**Supplementary Fig. 11b**). From the ribosome 3D reconstruction (**Supplementary Fig. 11a, Supplementary Fig. 11c** (left), **Supplementary Table 1)** 3D variability analysis^63^ was performed to further delineate ribosomal classes. Subsequent 3D reconstructions to resolve discrete heterogeneity across individual ribosomes reveal the presence of all major translation states with bound tRNAs at specific sites in the decoding center at resolutions from 10 to 13 Å (FSC=0.143, **Supplementary Fig. 11c, Supplementary Fig. 12a-f**), similar to the states derived from human ribosomes from multiple samples^64^. Overall, occupancies by tRNAs of the exit, peptidyl and aminoacyl sites are derived, as well as movements of the L1 stalk (**Supplementary Fig. 11c**). These results show that the L-A virus is present in a cell extract, in proximity to ribosomes captured in active translational states.

Identified L-A viruses within the cryo-EM micrographs, besides being in proximity to ribosomes and flexible macromolecules appeared pleomorphic (**Supplementary Fig. 10a, Supplementary Fig. 10c, Supplementary Fig. 13**). Their structural states resembled the states shown in **Fig. 1a** but statistical analysis of the native cell extract revealed that most particles identified are in their mature state (**Supplementary Fig. 10c (I, VIII, IX**), **Supplementary Fig. 10d-e**) while other rare pleomorphic states were present (**Supplementary Fig. 10c (II-VII), Supplementary Fig. 13**). Also, the possibility is high that indented particles were counted with the states due to shape similarity.

All observed particles are in proximity to ribosomes and irregularly shaped macromolecules, possibly mRNA, and may be involved in, *e.g*., mRNA decapping considering the L-A virus capsid function. To check if these particles are broken during cell lysis, fractions were followed by western blotting against the L-A virus gag (**Methods**). Results show a high abundance of the virus in the high molecular weight SEC fractions (**Supplementary Fig. 10f, Supplementary Fig. 14**), while, in the lower-molecular weight fractions, immunodetection of the viral capsid components did not yield a result (**Supplementary Fig. 10f, Supplementary Fig. 14**) implying that the states observed could correspond to particles with minimal damage stemming from biochemical treatment. In addition, the distribution of pleomorphic states and their non-random distribution towards a higher abundance of empty and full viruses corroborates the visualization of relatively unperturbed particles (**Supplementary Fig. 10d**). The packaging efficiency of the virus, *i.e*., mature viruses encapsidating the genome, defined by the ratio between randomly counted full (*N=400*) or empty (*N=661*) mature capsids is ~40% **(Supplementary Fig. 10e)**. At this resolution, we are unable to make any conclusions concerning the assembly mechanism^65^.

A final frequent observation in the cryo-EM micrographs of L-A virus-containing yeast native cell extracts was formation of small groups of the L-A virus (**Fig. 6a** and **Supplementary Fig. 15**). In these groups, composed of 12 ± 4 viruses on average, the L-A virus is mostly in a mature form and in proximity to other proteins and RNA, but with the L-A virus communities separated from other single particles in the images (**Fig. 6a**). To investigate this observation, we imaged high-pressure frozen and vitrified 30 nm thick whole yeast cell sections by transmission electron microscopy. These sections show similar groupings (on average 16 ± 10 viruses, **Fig. 6b**). Distances between the closest viruses and their measured diameter are comparable statistically (**Fig. 6c-d**), corroborating their in-extract and in-cell correspondence. Such assemblies have been widely reported for exogenous viruses and are frequently associated with either membranes or other cytoskeletal features, for example being connected to the Golgi apparatus or exploiting actin filaments^3^. In our cell section images, the L-A virus communities show no obvious interactions with the cytoskeleton or membranes but colocalize with the translation machinery, *i.e*., mostly ribosomes and irregular, electron-dense macromolecules that could represent mRNA (single-particle micrographs). These viral communities appear to be nano compartments that are relatively isolated from other cellular material.

**Figure 6.**
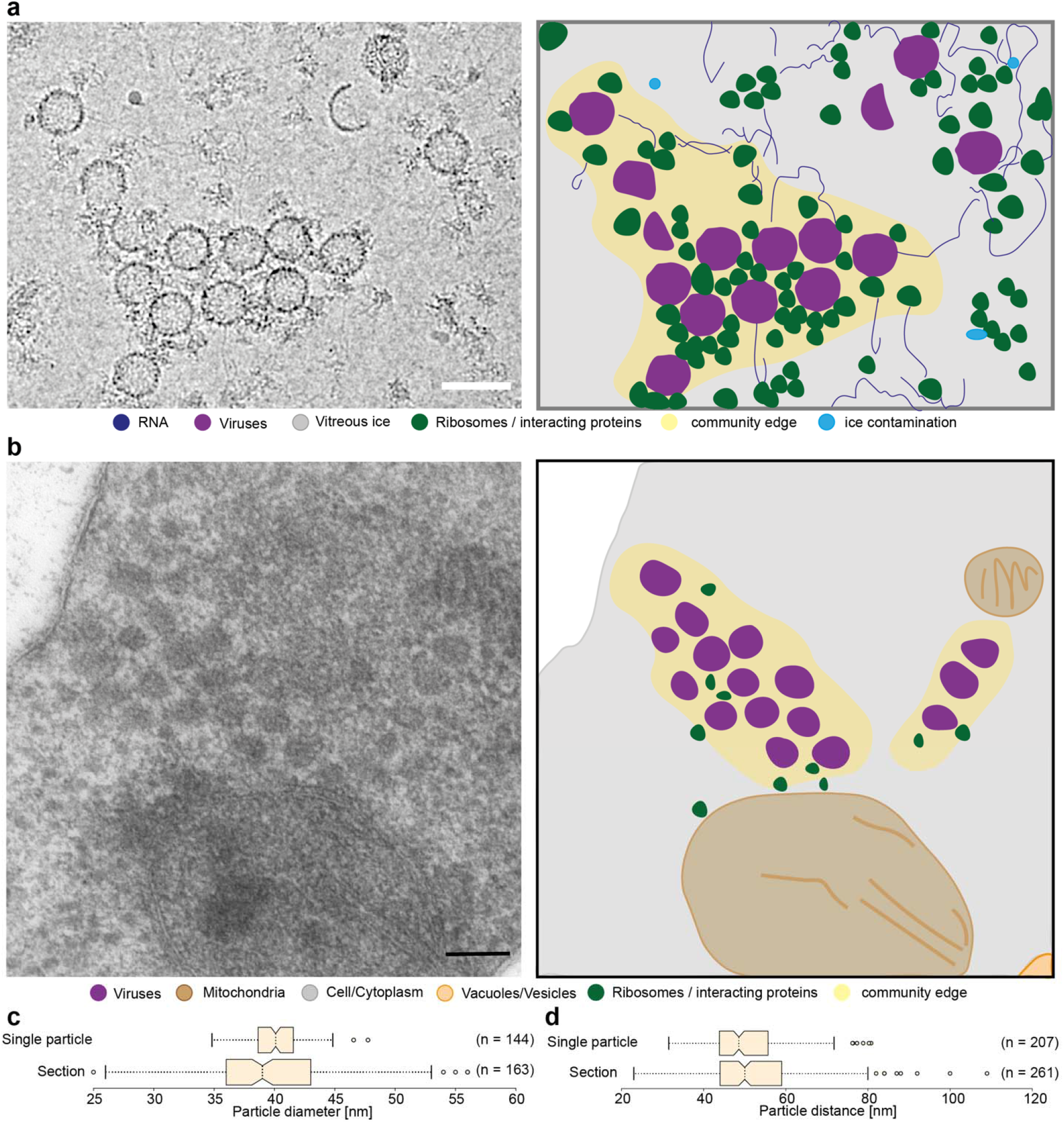
L-A virus communities in native cell extracts and in yeast cells: (a) Proposed viral communities are shown in a crop of a micrograph. Molecules are colored, such as RNA in dark blue, ribosomes in dark green, ice contamination in light blue and L-A virus in purple. The space around the group of viruses and surrounding proteins is colored yellow. (b) Cryo substituted, in resin embedded *Saccharomyces cerevisiae*, cut into sections of 30 nm in width and imaged on the EM 900. Shown is a crop through yeast on the left and an impression on the right. Color coding stays the same as in (a). The measured particle diameter (c), as well as the distances between L-A viruses (d), are represented by box plots with the standard deviation shown in the whiskers. All scale bars represent 50 nm.

## Discussion

Fast and reliable screening and analysis of viral particles in the context of their native assembly in eukaryotic cells is the prerequisite for the fast development of anti-viral treatment – a global need, as shown by a large number of viral infection events. Already in the 21^st^ century, multiple severe viral infections happened, *e.g*., SARS-Cov1 and -Cov2, multiple Ebola outbreaks, zoonotic influenced strains (swine flu, avian flu, etc.), and the Zika virus^66^. Here, we demonstrate a straightforward yet mild and “native-state-preserving” protocol for analyzing viral structures in their native context that is even applicable to endogenous viruses that are of low cellular abundance. While viruses develop numerous mechanisms to survive and replicate in the host cell, including hijacking cellular pathways^3^, nuclear import (e.g. HIV)^67^, and integration into the host genome^33^, the viral capsid formation is always the final step during maturation. Using the L-A virus as a model, we show that this does not happen at random cytoplasmic regions, but in communities that may involve different steps in capsid maturation, associated with the translation apparatus.

Our work also reveals critical insights into virus structure in the endogenous context: The reconstruction of the L-A viral capsid from native cell extract using cryo-EM attains a resolution comparable with the previously published, highly purified structure using x-ray crystallography^37^ and stems from a sample which is captured in a highly heterogeneous environment, interacting with other electron-dense cellular material (*e.g*., compare **Fig. 1c** to Figure 1 from *Castón et al*.^36^).

Unsurprisingly, the core regions of the capsid do not show any major differences, but the solvent-accessible flexible loop regions show clear deviations, which may represent functional differences in the native environment. Interestingly, multiple cation-π interactions are identified within the capsid subunits that are not only major contributors to the stability of the capsid but also may assist in folding and higher-order assembly. This is because a cation-π interaction needs the exact placement of both partners, *i.e*., the delocalized π-electron system and the cationic moiety. Once formed, this interaction is rigid and has clearly visible well-resolved side chains.

As previously described, the L-A capsid harbors a decapping and ‘cap-snatching’ activity^37,41,42^, located at the capsid surface. By this, the virus can (a) cap its own mRNA for increasing the half-life, and (b) create mRNA decoys for degradation. We identify a structured region at the active site, which shares structural homology with the methyltransferases and the colicin-E5 immunity protein. As the L-A virus decapping activity is specific only for methylated cap structures, a methyltransferase activity (or the capability of recruiting endogenous methyltransferases), might assist the endogenous activity, by methylating endogenous mRNA prior to decapping. Additionally, the E5 immunity protein is able to neutralize the colicin-E5, which is a tRNAse, by mimicking RNA.^37^ In the context of the L-A virus, this could serve as an interaction point for RNA binding proteins, which might degrade the viral mRNA, or by recruiting RNA-specific enzymes, *e.g*. methyltransferases, to the decapping site, again, optimizing intrinsic decapping activity.

Overall, by analyzing the L-A virus in its near-native environment, we were able to identify another layer of the viral architecture. By forming communities, the newly transcribed viral mRNA is immediately translated by the associated ribosomes, forming clearly visible polysomal structures at the site of capsid formation. By decorating the mRNA with a tight layer of ribosomes, mRNA degradation cannot take place. Putting the ‘cap-snatching’ mechanism in this context, capping the viral mRNA might not primarily serve as protection, but as an enhancer for translation of the viral proteins.

## Methods

### Cell cultivation and lysate fractionation

The *Saccharomyces cerevisiae* strain from ATCC (American Type Culture Collection PO Box 1549 Manassas, VA 20108 USA; ATCC^®^ 24657^™^) was cultivated in YPDG medium at 30°C for 5 h, to a OD595 of 2.5 (early exponential phase), then harvested at 3000 *g* for 5 min at 4°C and washed with distilled water (resulting pellet ~7 g). The resulting pellet was then lysed via bead beating using glass beads and lysis buffer (100 mM HEPES pH 7.4, 95 mM NaCl, 5 mM KCl, 1 mM MgCl2, 5% Glycerol, 0.5 mM EDTA, 1 mM DTT, 10 μg/ml DNAse, 2.5 mM Pefabloc, 40 μM E-64, 130 μM Bestatin, 0.5 μM Aprotinin, 1 μM Leupeptin, 60 μM Pepstatin A) in 3 rounds for 20 s of 6.5 mps in a fast prep. The lysate was centrifuged for 5 min at 4000 *g* at 4°C (Beckmann Heraeus multifuge 16R Swinging bucket rotor). The supernatant was again centrifuged at 100000 *g* for 45 min at 4°C (Beckman Optima^™^ MAX-XP Rotor MLS-50). After filtering the supernatant through a 0.22 μm syringe filter, the filtered lysate was concentrated using an Amicon with a cut-off of 100 kDa. Protein concentration was measured using Bradford reagent.

The lysate (concentration of 14 mg/ml) was loaded on a size exclusion column type Biosep 5 μm SEC-s4000 500 Å LC-Column 7.8×600. The system used was an ÄKTA Pure 25M FPLC; the fractionation was performed at a 0.15 ml/min flow rate, with a 200 mM NH_4_CH_3_COO-buffer pH 7.4 (filtered and sonicated), and fractions of a total volume of 250 μl were collected (**Fig. 1b**). The eluted fractions concentration was determined using Bradford reagent and directly used for data acquisition or flash-frozen and stored at −80°C until further use.

### Sample preparation for cryo-EM and data collection

To prepare samples for cryo-electron microscopy, holey carbon-coated copper grids (type R2/1 200 mesh from Quantifoil) are glow discharged using a PELCO easiGlow (15 mA, grid negative at 0.4 mbar, and 25 s glowing time). Directly after glow discharging the grids, 3.5 μl of the sample at a concentration of ~0.3 mg/ml is applied to each grid and plunge frozen using the Vitrobot^®^ Mark IV System (Thermo Fisher Scientific) and standard Vitrobot Filter Paper (Grade 595 ash-free Ø55/20 mm). Vitrification conditions were as follows: humidity 100%, temperature 4°C, blotting time 6 sec, blot force 0, sample volume 3.5 μl, blot offset 2 mm. After clipping, the vitrified grids were loaded onto a Thermo Fisher Scientific Glacios 200 kV Cryo-transmission electron microscope. The Falcon 3EC direct electron detector was used for acquisition. The images were acquired in linear mode with a total dose of 30 e^−^/Å^2^ (the acquisition parameters are shown in **Supplementary Table 1**).

### Image processing, analysis and model refinement

Two datasets were acquired for this work. One large dataset with 10 acquisitions of fractions 5 and 6 at a pixel size of 3.17 Å and a second smaller one with 2 replicates of fraction 5 with 1.57 Å. The 3.17 Å dataset was first analyzed with Scipion3^68–71^ and the refined map was then transferred to CryoSparc for further analysis. The 1.57 Å dataset was processed using CryoSparc only. The movies were firstly motion-corrected^72^ and then CTF^73^ estimation was performed before particle picking. Particle picking was performed manually and as a prerequisite for template-based picking, yielding approximately 230000 particles. For the dataset, at 1.57 Å pixel size, manual picking was omitted and template picking was directly performed yielding 2.7 million particles. Both datasets were subjected to sequential rounds of 2D and 3D classification. After removing low-resolution data, a final dataset of 65000 particles (3.17 Å pixel-size acquisition) and 17000 particles (1.57 Å pixel-size acquisition) was used for 3D reconstruction. The resulting maps were refined, by applying icosahedral symmetry and also using the option of symmetry expansion in CryoSparc2^74^. In the end, the FSC was calculated and the local resolution was derived according to the gold standard FSC^68^. The icosahedrally averaged map containing two independent protomers was of sufficient quality to build a Cα trace using Coot^75^. A subsequent DALI search^56^ identified the X-ray structure of the L-A virus (PDB ID: 1m1c) as best hit. Thereafter, the model was rigid-body fitted into the density using ChimeraX^76^ and subjected to iterative cycles of real space refinement using PHENIX^77^ and manual refinement using Coot to yield the final coordinates.

The DALI webserver^56^ was also used to identify structural homologues of the cap-snatching domain. As input structure, the region 491-587 from chain A was used. To identify suitable hits, the “Matches against full PDB” was used, and a scoring threshold was set to Z > 3.0, which is an optimized similarity score between two structures, in which the higher the number the greater the similarity.^78^

### Energetic calculations

The guru- and multi-body-interface of the HADDOCK webserver 2.2 ^55,79^ was used to calculate protein-protein-interaction energetics. All clustered models were used to calculate the energetic distribution. Calculations were done using an inhouse python script. For calculations of capsid stability, the capsid structure was used as a template and, starting from a monomeric subunit, the energetics for each possible monomeric interaction partner was calculated, using the 9 unique interfaces of the capsomere (see **Supplementary Fig. 5**), and the formula:

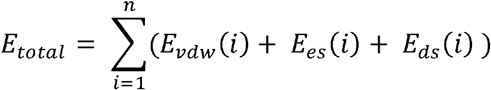

Best ranked “docked” monomer were selected. This was repeated, till the entire capsid was built. Calculations were done by a PyMOL python script.

### Calculation of map cross-correlation coefficients and model prediction with AlphaFold2

The PHENIX tool “Comprehensive validation (cryo-EM)” was used with default parameters^80^ to calculate map cross-correlation coefficients. The Gag-Pol fusion protein of the *Saccharomyces cerevisiae* L-A helper virus was predicted by the in-house installed AlphaFold2 (v.2.1) while using default parameters and the sequence as fasta-file as input.^58^

### Mass spectrometric analysis

In-solution digestion was performed for sample preparation for mass spectrometric (MS) analysis. 10 μg of fraction 5 or 6 (technical duplicates of biological triplicates) were precipitated and adjusted to 100 μl with ice-cold acetone and incubated for 60 min at −20 °C. Precipitated proteins were centrifuged (10 min, 20,000 *g*) and airdried. Protein pellets were resuspended in 25 μl of 8 M urea in 0.4 M ammonium bicarbonate, reduced with 5 μl of 45 mM DTT (30 min at 50 °C), and alkylated with 5 μl of 100 mM 2-chloroacetamide (30 min at 37 °C). The volume of each sample was adjusted to 200 μl by water. The samples were digested using trypsin (Promega Sequencing Grade Modified Trypsin) using a 1:50 (w/w) enzyme:protein ratio for 16 hours at 37 °C. The reactions were stopped by adding 10 μl of 10% (v/v) TFA. 20 μl of digestion mixtures were analyzed by LC/MS/MS using a U3000 nano-HPLC system coupled to a Q-Exactive Plus mass spectrometer (Thermo Fisher Scientific). Peptides were separated on reversed phase C18 columns (trapping column: Acclaim PepMap 100, 300 μm × 5 mm, 5μm, 100 Å, Thermo Fisher Scientific; separation column: μPAC 50 cm C18, Pharmafluidics). After desalting the samples on the trapping column, peptides were eluted and separated using a linear gradient ranging from 3% to 35% B (solvent A: 0.1% (v/v) formic acid in water, solvent B: 0.08% (v/v) formic acid in acetonitrile) with a constant flow rate of 300 nl/min over 180 min. Data were acquired in data-dependent MS/MS mode with higher-energy collision-induced dissociation (HCD), and the normalized collision energy was set to 28%. Each high-resolution full scan (*m/z* 375 to 1799, R = 140000 at *m/z* 200) in the orbitrap was followed by high-resolution fragment ion scans (R = 17500) of the 10 most intense signals in the full-scan mass spectrum (isolation window 2 Th); the target value of the automated gain control was set to 3,000,000 (MS) and 200,000 (MS/MS), maximum accumulation times were set to 50 ms (MS) and 120 ms (MS/MS). Precursor ions with charge states <2+ and >6+ or were excluded from fragmentation. Dynamic exclusion was enabled (duration 60 seconds, window 3 ppm).

### Identification of most abundant proteins from MS/MS data

MS-raw files were analyzed with MaxQuant (version 1.6), with activated label-free quantification (LFQ), iBAQ and “Match between runs” option, and the annotated yeast proteome (UP000002311). For identification of viral proteins, the sequences of the following L-A virus proteins were added to the database: gag (UniProt-id: P32503) and pol (UniProt-id: Q87022; residue-range: 647-1505), the L-BC gag-pol (UniProt-id: P23172), the ScV-M1 preprotoxin (UniProt-id: P01546), the ScV-M2 preprotoxin (UniProt-id: Q87020), and the ScV-M28-like killer preprotoxin (UniProt-id: Q7LZU3). MaxQuant derived results files were analyzed and plotted by an in-house python script.

### Western Blot analysis

Gels were freshly casted in-house prior to the experiment using a separating gel: 10% (w/v) acrylamide (37.5:1), 0.1% (w/v) SDS, 0.04% (w/v) APS, 0.002% (w/v) TEMED in 370mM Tris-HCl solution pH8.8 and stacking gel: 5% (w/v) acrylamide (37.5:1), 0.1% (w/v) sodium dodecyl sulfate (SDS), 0.04% (w/v) APS, 0.002% (v/v) TEMED in 125□mM Tris-HCl-solution pH 6.8. The gels cast were 1mm thick. The samples were mixed with a 4x loading dye (250□mM Tris-HCl (pH 6.8), 8% w/v SDS, 0.2% w/v bromophenol blue, 40% v/v glycerol, 20% v/v β-mercaptoethanol) and incubated for 5 min at 95°C shaking. Roughly 3 μg sample of the fractions and 2.5 μg as well as 5μg of the positive control, was loaded and electrophorized with a standard of 5□μl of Precision Plus Protein^™^ All Blue Prestained Protein Standards (BioRad #1610373). For electrophoresis, a 1x electrophoresis buffer freshly prepared using a 10x stock solution (30.3□g Tris-base, 144□g Glycine in 1□L of deionized water) was used and the gels run at an electrical field of 100□V for ~2□h. The gels were then transferred onto a nitrocellulose membrane using the Trans-Blot^®^ Turbo^™^ Transfer System of BioRad. A pre-set protocol of 25□V (1□A) applied field, for 30 min was used. After blotting the membranes were blocked using 5% (w/v) skimmed milk powder in TBST for 1h at 4°C. The primary antibody (rabbit anti-ScV-L-A peptide serum by GenScript) was diluted at 2:25000 in 2% (w/v) skimmed milk powder in TBST and incubated for 16h at 4°C. After incubation, the membrane was washed 3 times for 10 min with 2% (w/v) skimmed milk powder in TBST. The secondary antibody (goat anti-rabbit IgG H&L (HRP) ab205718 by abcam) was prepared the same way as the primary but at a final concentration of 1:25000. The membranes were incubated for 1h at RT. After another 3 washing steps, the membrane could be imaged using the ChemiDoc MP Imaging system and a freshly prepared ECL fluorescent mixture. Antibodies were custom-made by GenScript (New Jersey, USA), using the sequences provided in the Supplementary source data.

### High-pressure freezing

Yeast cells (harvested as described in the section Cell cultivation and lysate fractionation) were rapidly frozen with a high-pressure freeze fixation apparatus (HPM 010; BALTEC, Balzers, Liechtenstein). The material was cryo substituted with 0.25% glutaraldehyde (Sigma) and 0.1% uranyl acetate (Chemapol, Prague, Czech Republic) in acetone for two days at −80°C using cryo substitution equipment (FSU; BAL-TEC) and embedded in HM20 (Polysciences Europe) at −20°C. After polymerization, samples were cut with an ultramicrotome (Ultracut S, Leica, Wetzlar, Germany). The ultrathin sections (30 nm) were transferred to formvar-coated copper grids and poststained with uranyl acetate and lead citrate in an EM-Stain apparatus (Leica) and subsequently observed with a Zeiss EM 900 transmission electron microscope (Carl Zeiss Microscopy GmbH, Jena, Germany) operating at 80 kV.

### Reporting Summary

Further information on experimental designs can be found in the associated supplementary materials linked to this manuscript.

## Supporting information

04 Supplemental Validation ScVLA 3.77

05 Supplemental Validation ScVLA 6.4

06 Supplemental Validation Ribosome

## Data availability

The 3D maps of the L-A virus at 3.77 Å and inner density C1 reconstruction (EMD-15214) and the in-extract ribosome (EMD-15215) will be available in the EMDB database. The high-resolution map (EMD-15189) and the atomic model of yeast L-A virus (PDB-8A5T) are deposited in the EMDB and PDB, respectively. The proteomics data set can be accessed through PRIDE (Identifier: PXD034431). Energetic calculations and AlphaFold models are available at the SB-Grid database under the accession code 930. Source data are provided with this paper and all other data for support of this study are available from the corresponding author upon request. Primary cryo-EM movies will be deposited in EMPIAR (Accession code EMPIAR-11069).

## Code availability

All unpublished code and scripts used in this study are available upon request.

## Acknowledgements

The authors thank all members of the Kastritis laboratory for valuable insights, in particular Kevin Janson for growing yeast and Ioannis Skalidis for initially setting up data analysis of cryo-EM acquired fractions. The authors also thank Dr. Marta Fratini, and Prof. Dr. Ingo Heilmann (MLU, Halle, DE) for confocal microscopy of yeast protoplasts, as well as Prof. Dr. Stephan Feller and Dr. Lolita Piersimoni for valuable discussions. This work was supported by the Federal Ministry for Education and Research (BMBF, ZIK program) (Grant nos. 03Z22HI2 and 03Z22HN23 to PLK and 03COV04 to PLK and MTS), the European Regional Development Funds for Saxony-Anhalt (grant no. EFRE: ZS/2016/04/78115 to PLK and MTS), funding by Deutsche Forschungsgemeinschaft (DFG) (project number 391498659, RTG 2467), and the Martin-Luther University of Halle-Wittenberg.

## Author Contributions

L.S. performed sample preparation, cryo-EM grid preparation and screening, and F.L.K. supervised the laboratory work. F.H. collected the data. Data analysis and structure calculations were performed by L.S. under supervision from D.A.S. and P.L.K. M.T.S. build the initial model and identified the L-A virus via the DALI search. Structural modeling and structure validation was performed by C.T. MS analysis was performed by L.S. and C.T. C.I. measured MS data using A.S. instruments. G.H. performed cryo substitution experiments. A.M., P.N.M.S. and D.I.S. contributed to the conceptualization of the project. P.L.K. wrote the paper, with contributions from L.S., C.T. and all authors; L.S., C.T. and P.L.K. made the figures. P.L.K. conceived, funded, and supervised the project.

## Corresponding author

Correspondence to Panagiotis L. Kastritis

## Ethics declarations

*Competing interests:* The authors declare no competing interests.

## Notes

### Competing Interest Statement

The authors have declared no competing interest.

