## Supplementary material for "Delineating organizational principles of the endogenous L-A virus by cryo-EM and computational analysis of native cell extracts": 02 Supplemental Figures

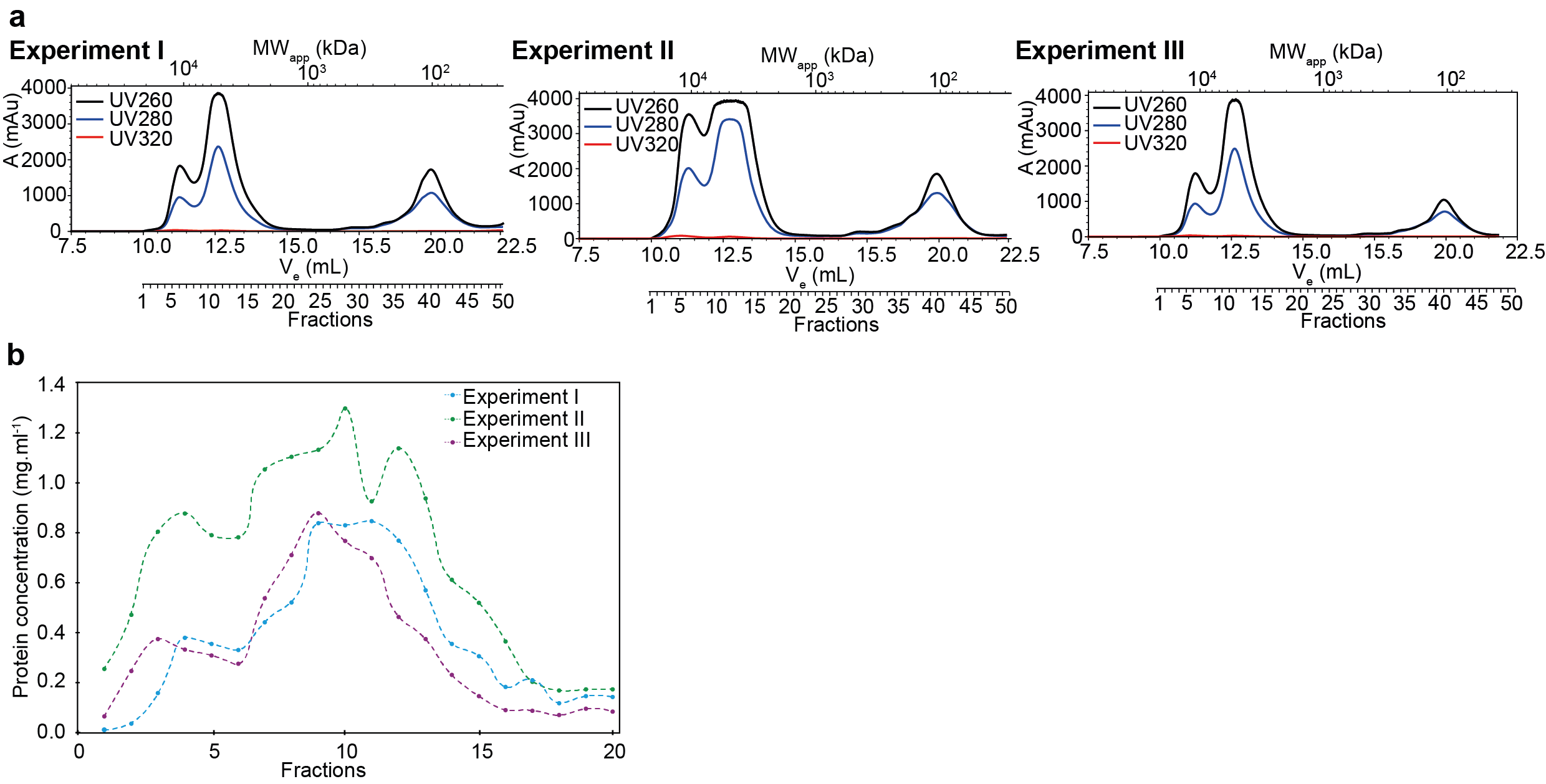

**Supplementary Fig. 1 Identification of the L-A virus:** (a) Triplicate SEC at 260 nm, 280 nm, and 320 nm absorbance. (b) Triplicate of the experiment showing high in-fraction protein concentration of the resulting fractions after SEC measured with Bradford reagent.

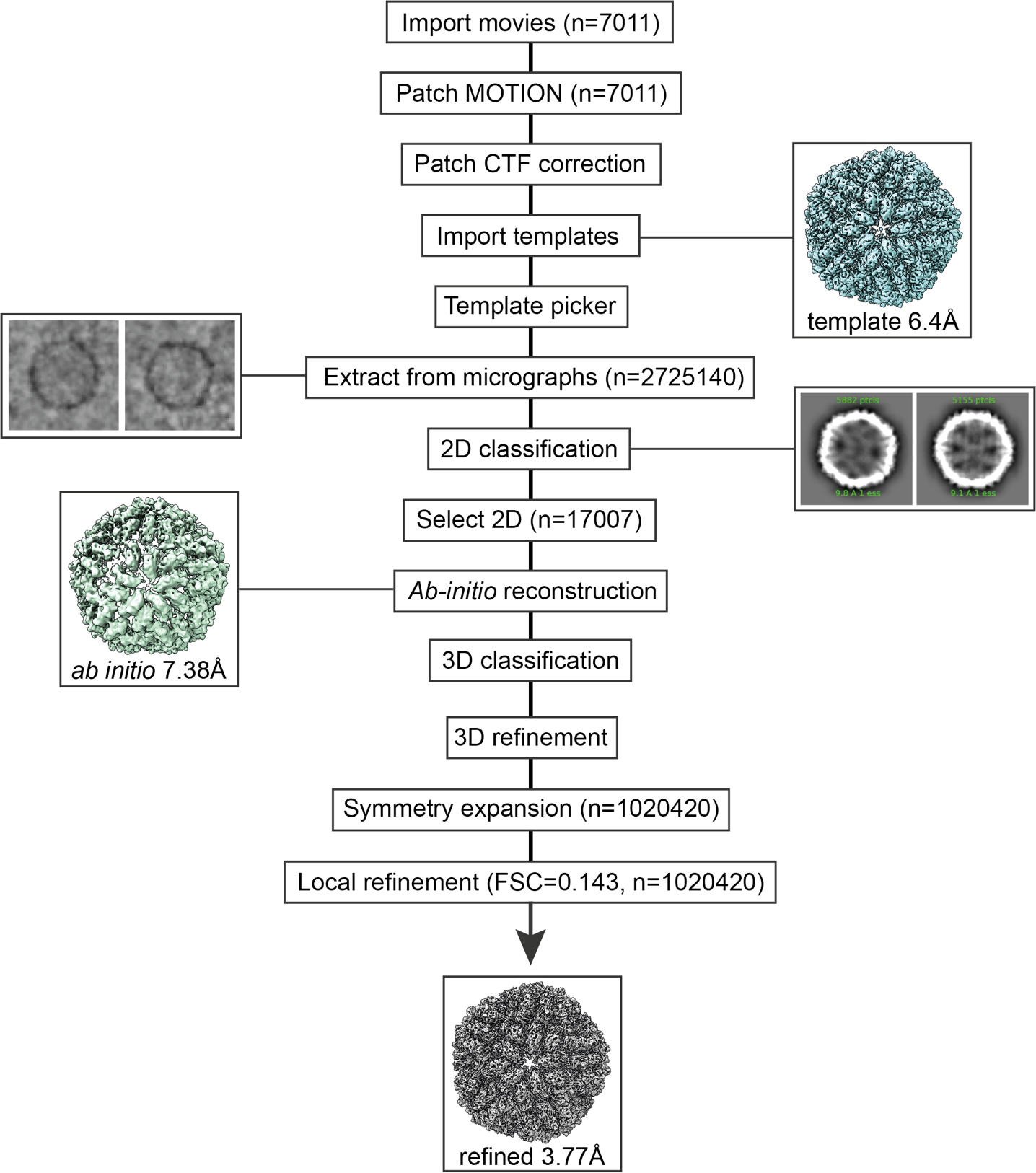
 **Supplementary Fig. 2 Workflow of image analysis:** Shown is a workflow of image analysis, specifically followed for the high-resolution cryo-EM structure of the L-A virus. Data is imported into the workspace, followed by motion correction and CTF correction procedures. After these processes finished, particles can be picked, and a *de novo* lower resolution reconstruction is derived after 3D classification and refinement. For template picking this reconstruction was chosen. The picked particles are then extracted and 2D classified. Selected 2D class averages were used for another *ab initio* reconstruction, followed by 3D classification and 3D refinement of chosen particles. The map quality was improved after using symmetry expansion and particles were, lastly, locally refined.

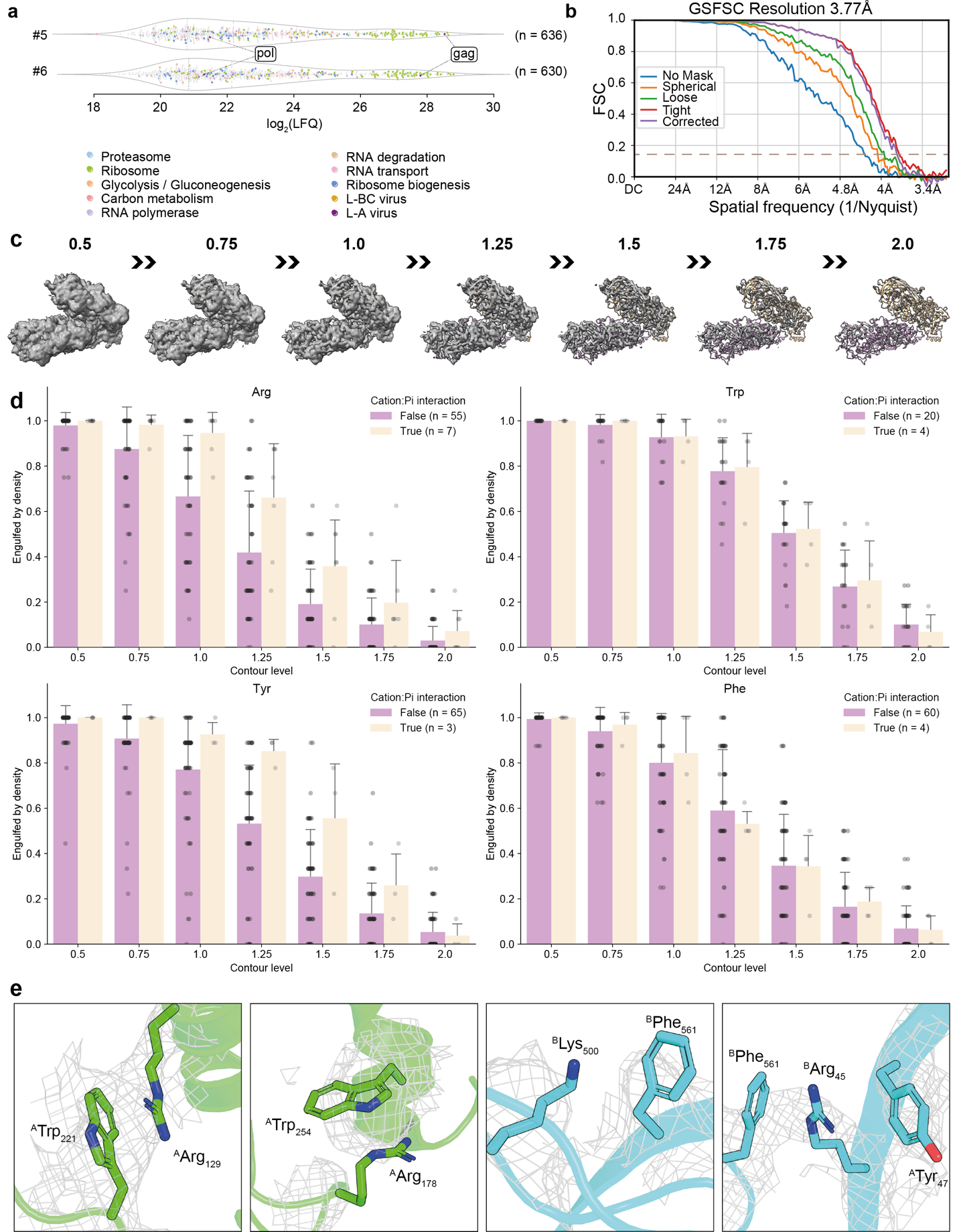
 **Supplementary Fig. 3 Mass spectrometry data for in-fraction protein content, cryo-EM resulting model and stabilizing interactions:** (a) Label-free quantification of the protein content of fractions 5 and 6 showing the virus proteins, translation-related proteins, proteins for glycolysis, carbon metabolism, spliceosome, citrate cycle, and others, present in the same cellular fraction. (b) FSC of the cryo-EM derived map for the L-A virus utilizing icosahedral symmetry. (c) Asymmetric dimer of the yeast L-A virus model superimposed on the capsid densities at different thresholds (increase in steps of 0.25 sigma) set in ChimeraX. (d) Bar plot of the resolvability of residues involved in cation-π interactions (papaya whip) against residues that are not involved in such interactions (purple) (e) Cation-π interactions shown between arginine/lysine and tyrosine, tryptophan or phenylalanine within chain A (green), and within chain B (blue).

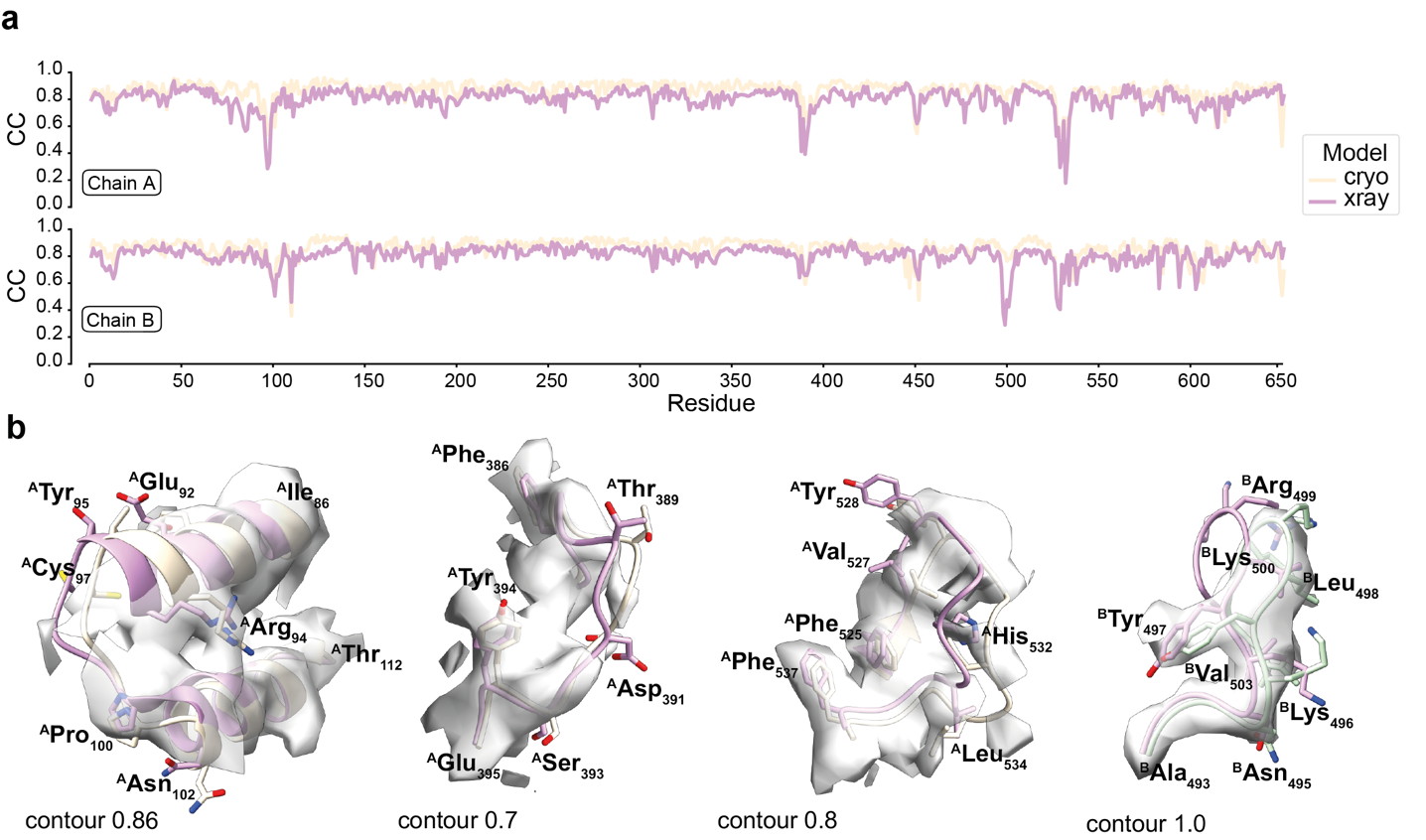
 **Supplementary Fig. 4 Fits and conformational variations of the X-ray and cryo-EM models to the resolved cryo-EM map:** (a) Cross-correlation of the cryo-EM (peach) and the X-ray (purple) models of chain A and chain B are shown in this plot, indicating local conformational variability as dips in cross-correlation values. (b) The cryo-EM- (peach) and X-ray-(purple) derived asymmetric homodimer fits for backbone (CC_cryo-EM_=0.85 vs. CC_X-ray_=0.79) and side-chain (CC_cryo-EM_=0.84 vs. CC_X-ray_=0.79) conformations. In addition, a comparison of cryo-EM (beige) and X-ray (purple) models is shown after fitting in the reconstructed capsid density (grey), highlighting locally substantial conformational differences.

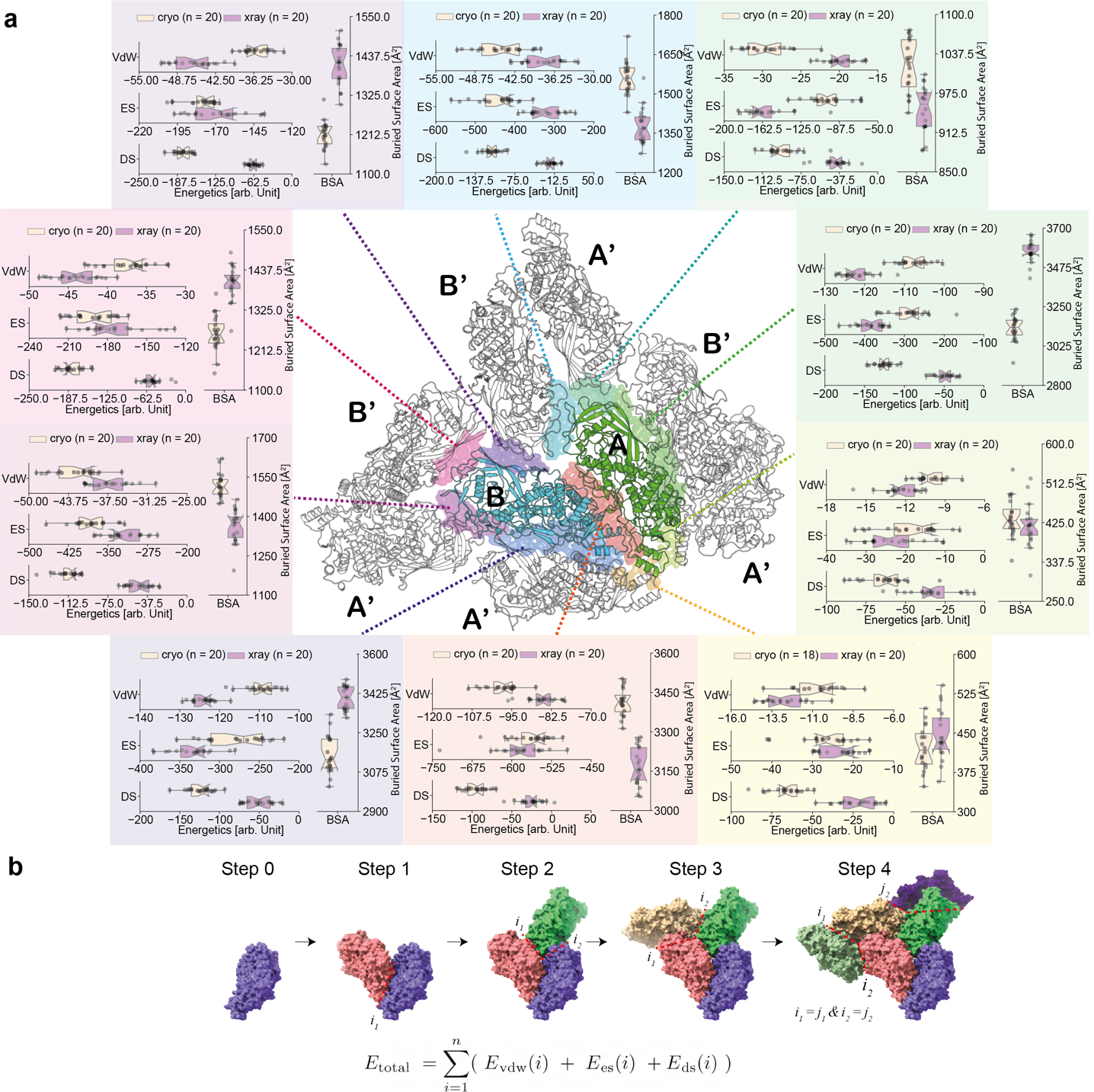
 **Supplementary Fig. 5 Comparison of interface energetics forming the crystallographically-resolved and cryo-EM resolved L-A virus capsid:** (a) Shown in peach (chain A) and green (chain B) in comic representation is the yeast L-A virus capsomer with adjacent protomers. Subunits corresponding to chain A are labelled with A’ and subunits corresponding to B are labelled B’. The colors for the calculated interfaces match the box-plot background color. Calculated were van der Waals forces (VdW), desolvation scores (DE), electrostatics scores (ES) in arbitrary units (a.u.), and the buried surface area (BSA) in Å^2^. (b) Proposed stability analysis (from monomer until step 4) of the capsid informed by the energetics calculations for derived interfaces.

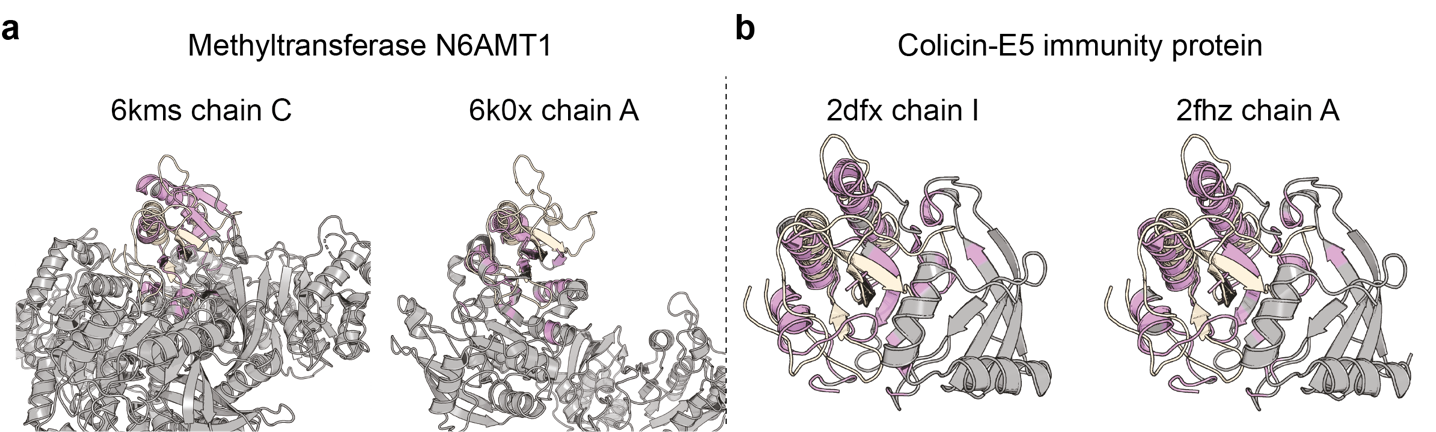
 **Supplementary Fig. 6 Identification of structural homologs using the DALI server:** Identified homologous domains of (a) the methyltransferase enzyme family and (b) the Colicin immunity protein family, both aligned to the mRNA decapping active site (residues I491-V587).

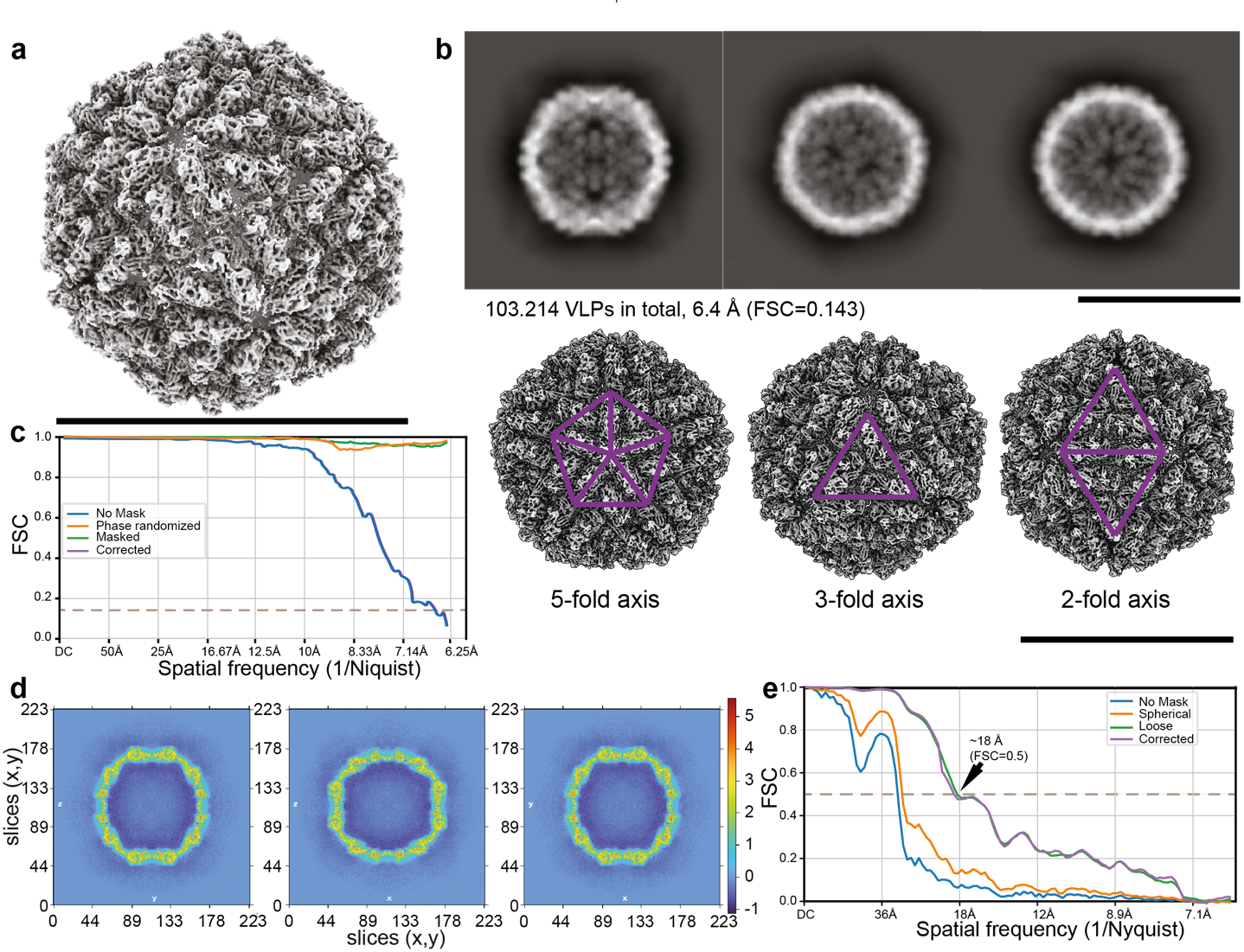
 **Supplementary Fig. 7 Inner view for the L-A virus:** (a) 3D reconstructed map of the L-A virus at 6.4 Å. (b) 2D class averages covering views of the 2-, 3- and 5- fold axes. (c) FSC of the 6.4 Å reaching Nyquist. (d) A high signal-to-noise ratio is achieved as shown by the map x,y slices. (e) FSC at 0.5 of the asymmetrically reconstructed inner densities after particle subtraction. Scale bars at 45 nm.

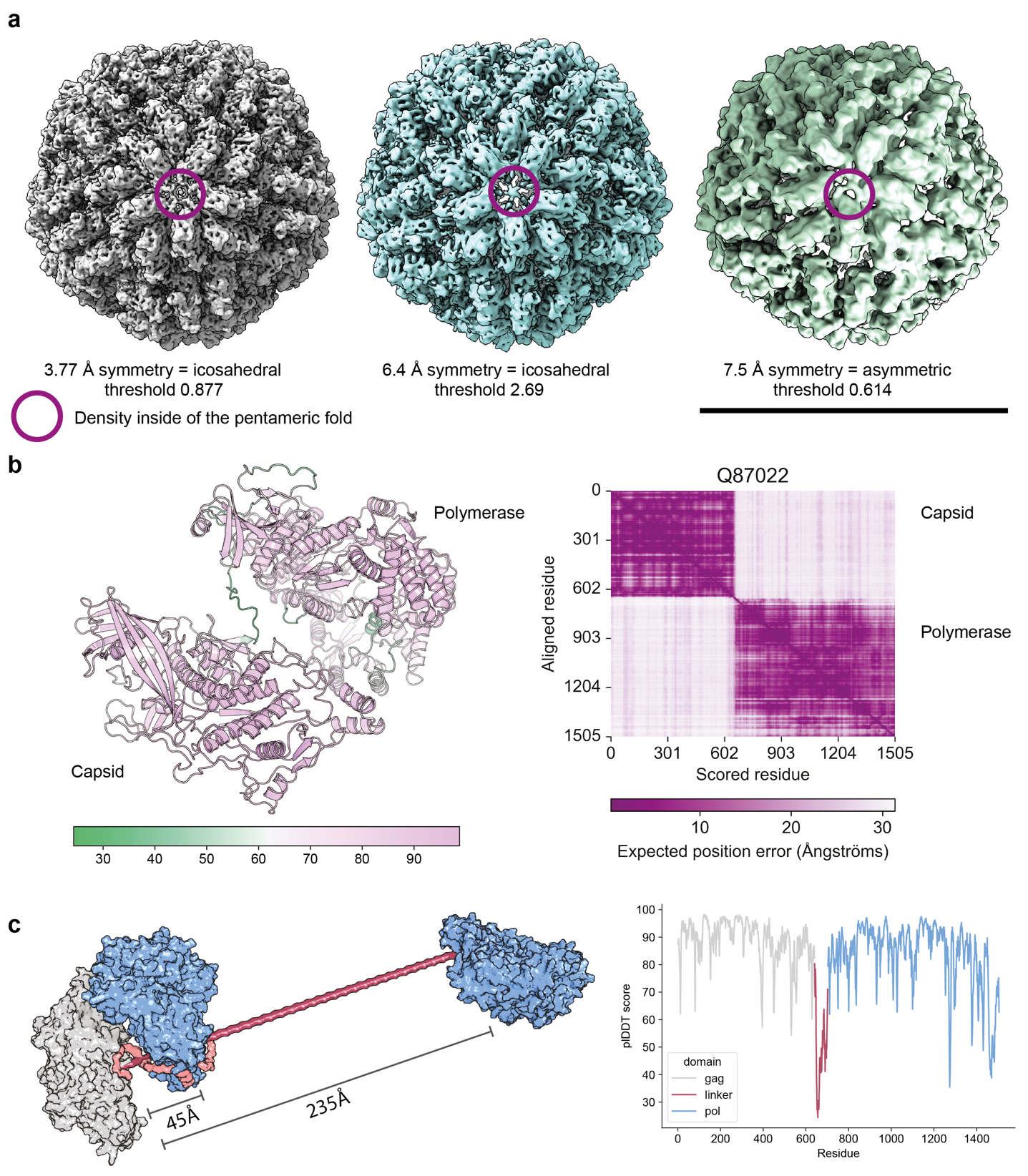
**Supplementary Fig. 8 Capsid-RNA polymerase communication:** (a) Cryo-EM maps of the Capsid showing the persistent density at the 5-fold axis. (b) AlphaFold2-predicted Gag-Pol fusion protein. The predicted structure of the polymerase is connected via a 62-residue flexible linker to the capsid structure. The heat bar below shows the confidence of AlphaFold2 for the model, the heatmap in purple shows the expected position error of each residue, while dark purple represents a low error and lighter color higher errors. The linker region (purple) shown in (c) is highly positively charged and supposedly positions the Pol (blue) relative to the Gag (grey). AlphaFold2-predicted and maximum distances are drawn at 45 Å and 235 Å, respectively. On the right, the pLlDDT score shows the low confidence prediction for the flexible linker region.

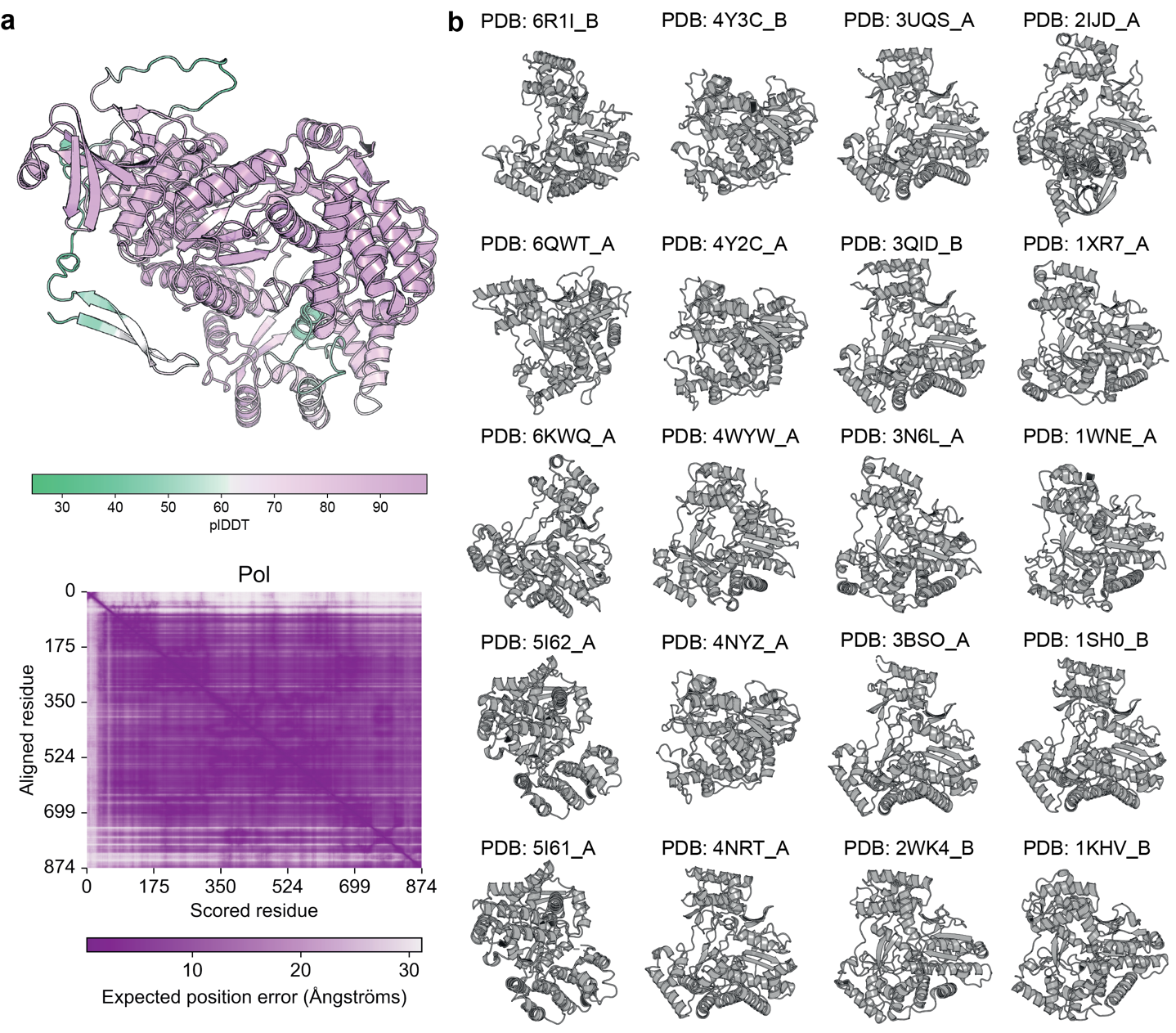
**Supplementary Fig. 9 Analysis of the RNA pol AlphaFold model:** (a) AlphaFold predicted polymerase protein. The heat bar below shows the confidence of AlphaFold for the model it created, the heatmap in purple shows the expected position error of each residue while dark purple represents a low error and lighter color high errors. (b) Shown are the templates AlphaFold used for predicting the yeast L-A helper virus polymerase model.

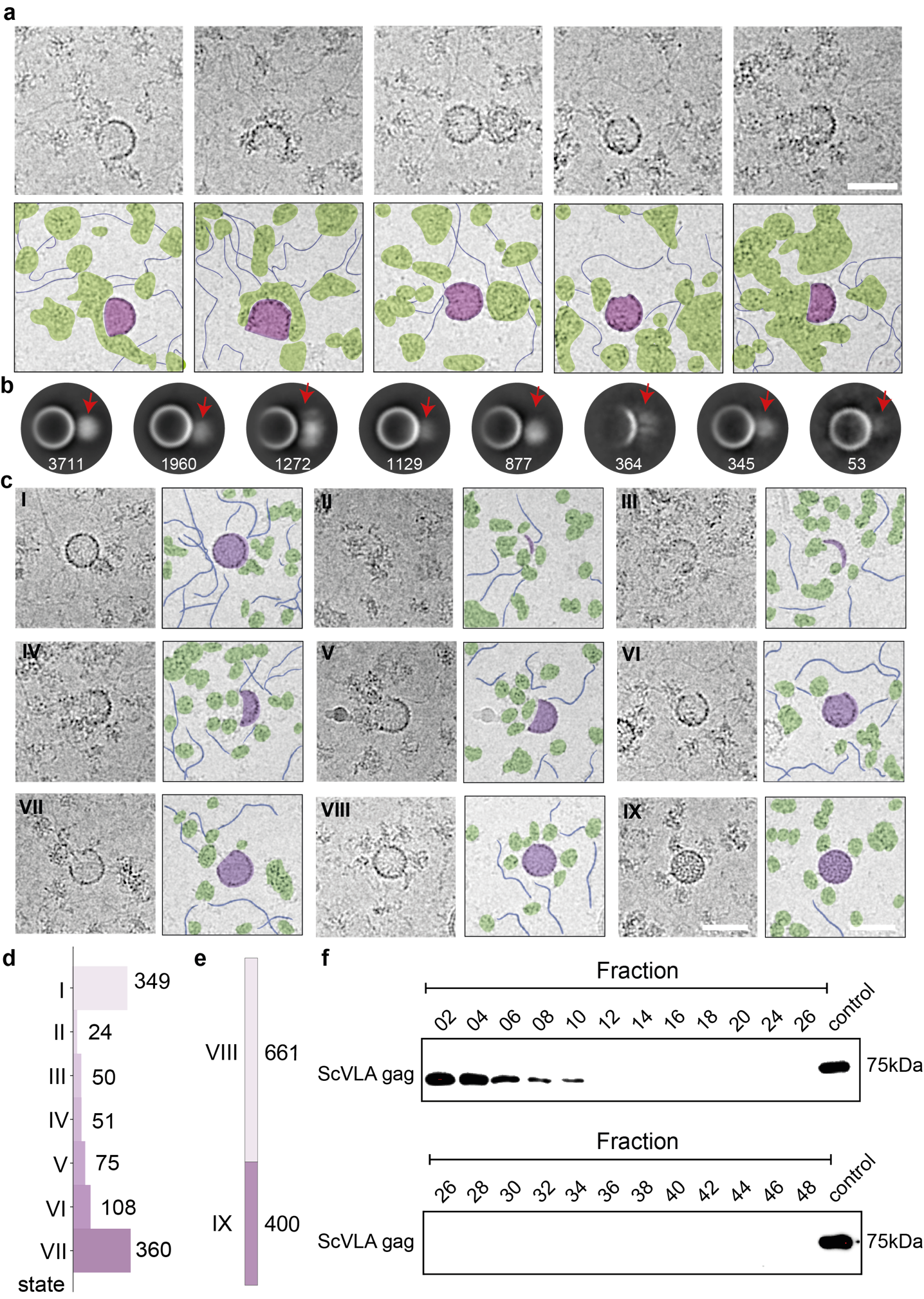
 **Supplementary Fig. 10 L-A virus environment and assembly:** (a) A representation of assembly states corresponding to the viral lifecycle is shown in crops and impressions in distinct steps from I to IX, like **Fig. 1a**. The scale bar is 50 nm. (b) 2D classes of the LA virus capsid with diffused densities in proximity; Ribosomes are often bound as shown in, *e.g.*, panel (a). (d) A statistical representation of the assembly states I to VII in absolute numbers. (e) Statistical comparison of full and empty viruses. This shows the ratio between randomly counted full (*N=400*) or empty (*N=661*) mature capsids corresponding to states VIII and IX of (c). (f) Western blot analysis against gag shows signal only in high molecular weight fractions indicating that the virus capsids are not likely to be severely damaged.

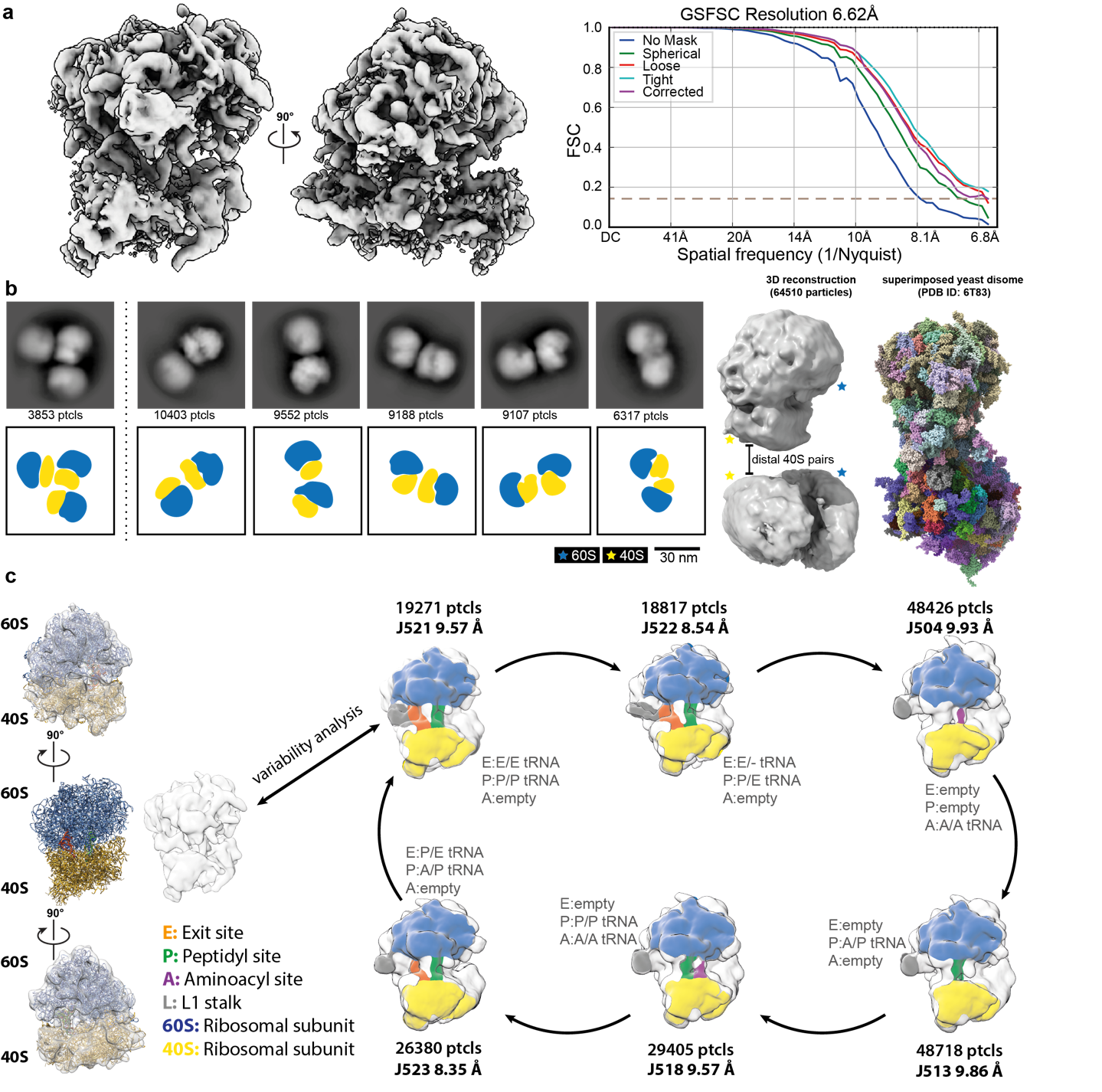
**Supplementary Fig. 11 Cryo-EM reconstructions of translationally active ribosomes:** (a) 3D reconstructed ribosome at 6.62 Å with an FSC=0.143. (b) 2D class averages and 3D reconstruction of identified polysomes within the native cell extract. Scale bar represents 30 nm. (c) Identified distinct translational states after variability analysis performed in CRYOSPARC.

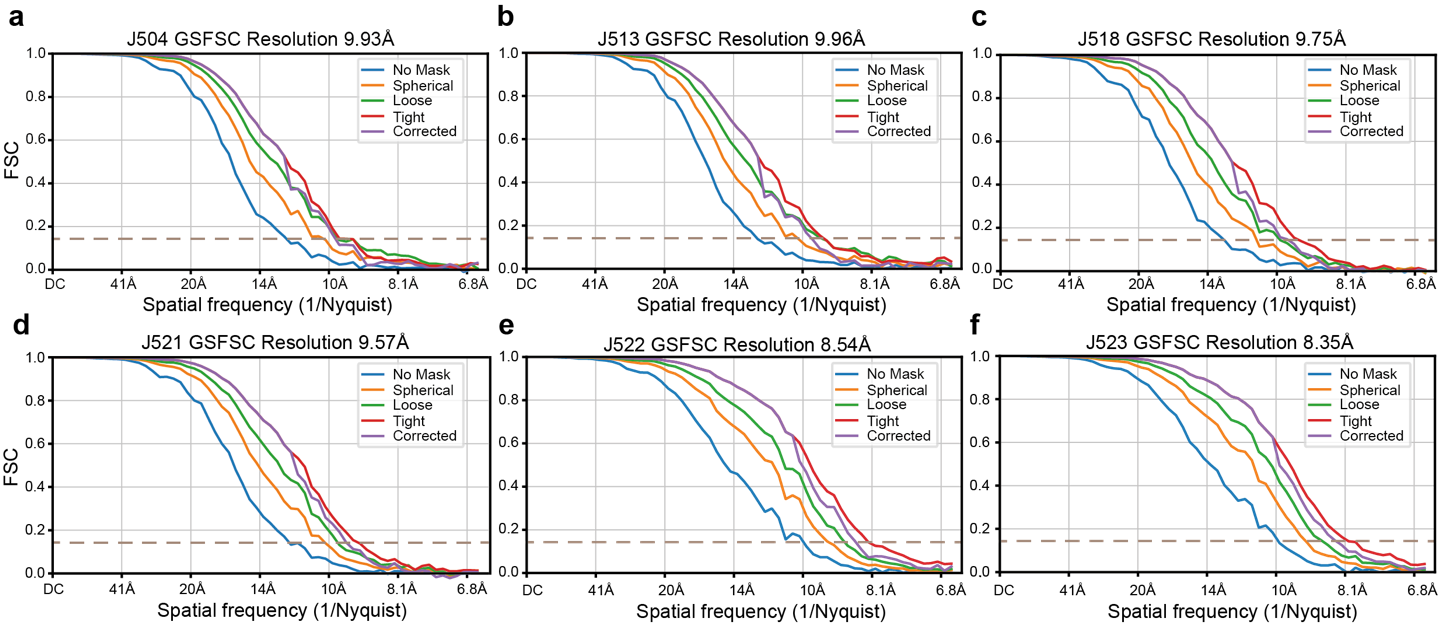
 **Supplementary Fig. 12** (a-f) FSCs of the 6 distinct translational stages of reconstructed ribosomes after variability analysis shown in **Supplementary Fig. 11c**.

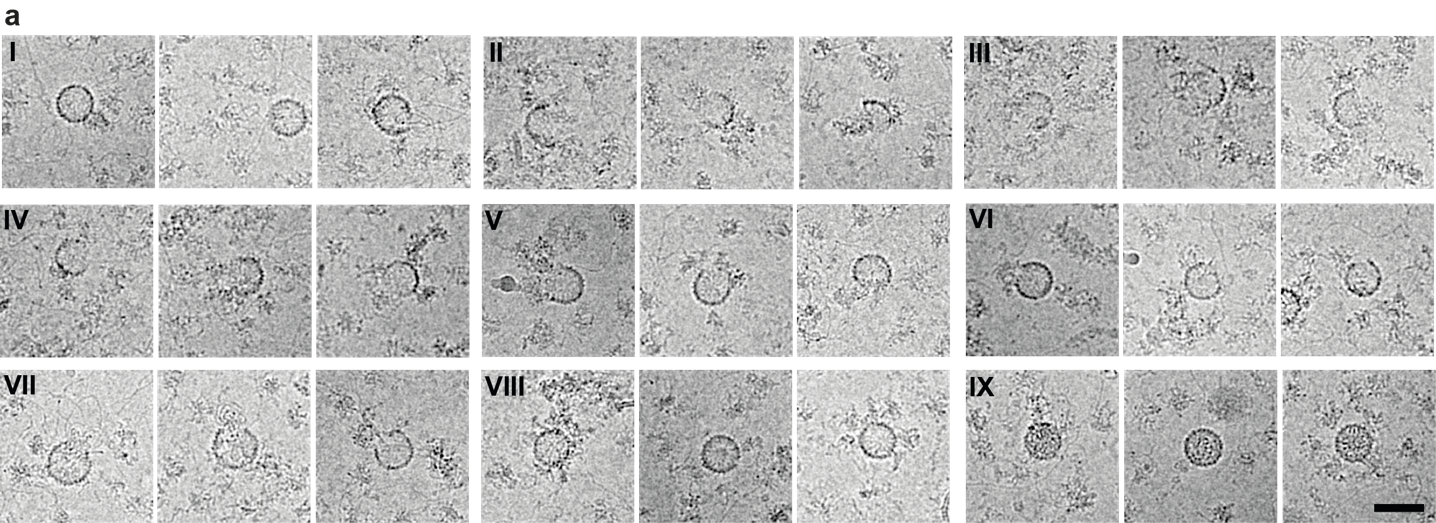
 **Supplementary Fig. 13 Potential viral life cycle states:** (a) More representations of the different states of the viral lifecycle shown in crops, distinctive steps from I to IX. The scale bar represents 50 nm.

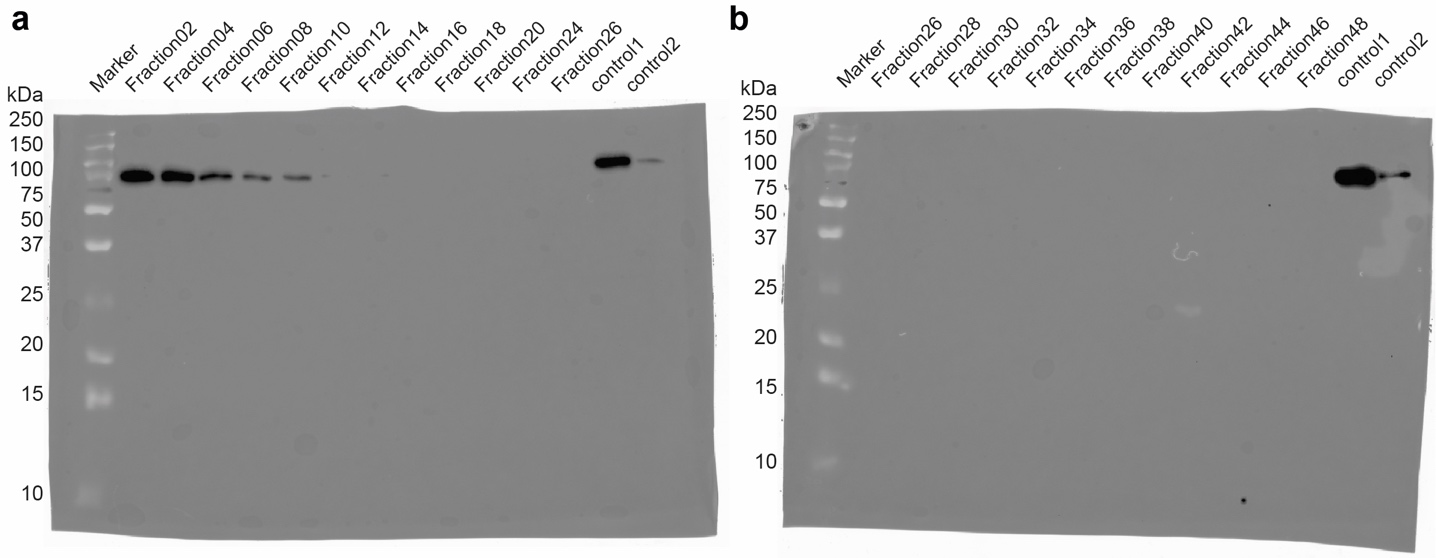
**Supplementary Fig. 14 Tracking of the abundance of the L-A helper virus via Western blot analysis:** (a, b) Original uncropped membranes after Western blot analysis. Every second fraction was applied on the gel and L-A virus capsid was detected. Control 1 and 2 are positive controls and correspond to the fraction in which the LA virus was detected by mass spectrometry (1), and the yeast lysate used for injecting the SEC (2).

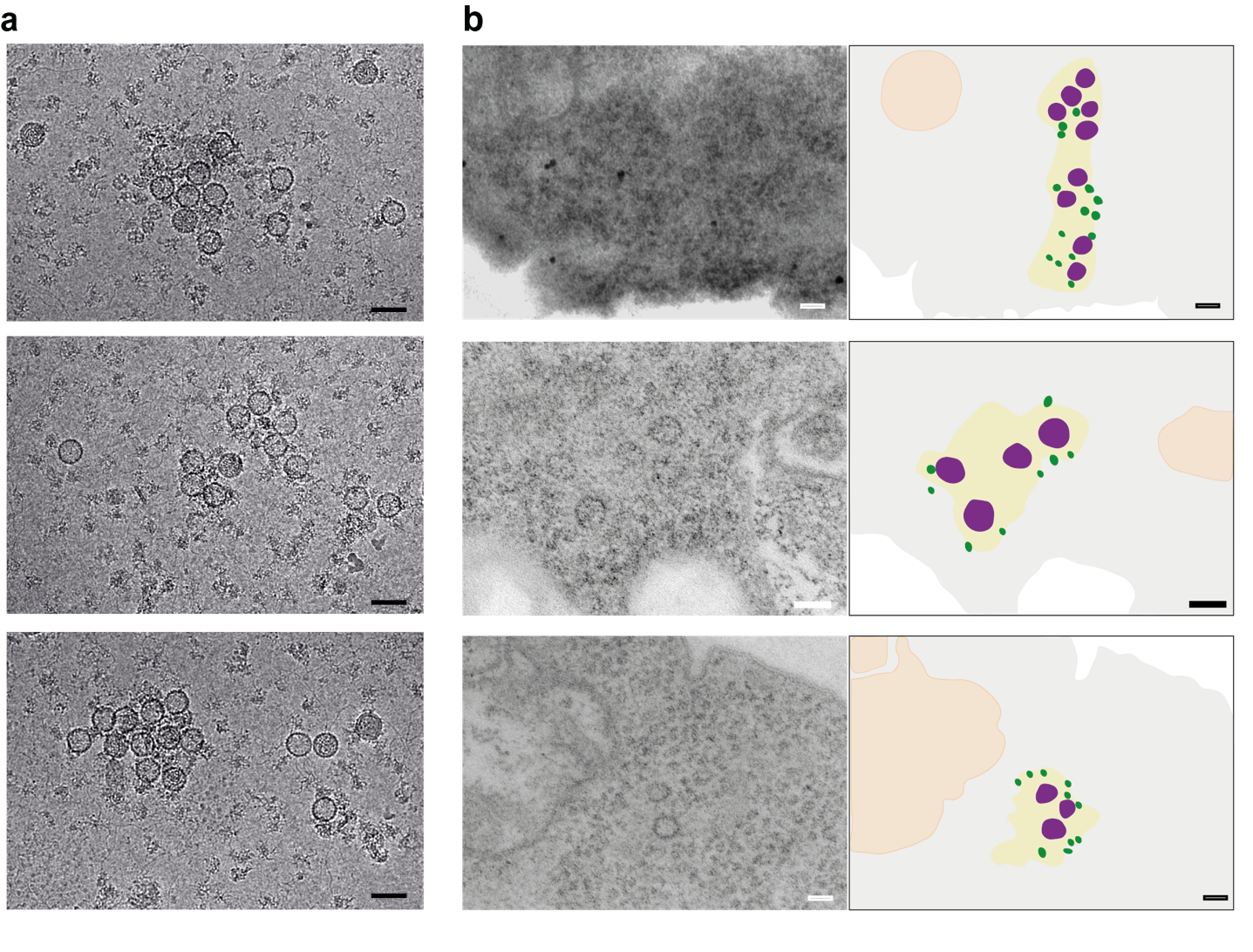

**Supplementary Fig. 15 L-A virus communities:** (a) More examples of viral particles grouping in the micrographs. Cryo-substituted, in resin embedded *Saccharomyces cerevisiae* sections of 30 nm in width and imaged on the EM 900. Shown are crops through yeast on the left and a comic representation on the right. Vesicles are colored in orange, ribosomes/adjacent proteins dark green, and viral particles in purple (b). Scale bar representing 50 nm.

**Supplementary Table 1a** related to Figures 4, 5 & 6: Cryo-EM acquisition and reconstruction parameters of specimen reported in this study.

|  | Acquisition 1  (L-A virus)  (EMD-15214) | Acquisition 2  (L-A virus)  (EMD-15189)  (PDB-8A5T) | Acquisition 3  (Ribosome*^1^)  (EMD-15215) | Acquisition 3  (Inner density L-A virus) |
| --- | --- | --- | --- | --- |
| **Data collection and processing** |  |  |  |  |
| Magnification | 45000X | 92000X | 45000X  200  TFS Glacios  TFS Falcon IIIEC  12*  30  2.3  -1.0 to -3.0  3.17  2728  TFS EPU 2  C1 |  |
| Voltage (kV) | 200 | 200 |  |  |
| Microscope model | TFS Glacios | TFS Glacios |  |  |
| Camera model | TFS Falcon IIIEC | TFS Falcon IIIEC |  |  |
| Number of frames | 12* | 12* |  |  |
| Electron exposure (e–/Å^2^) | 30 | 30 |  |  |
| Per-frame exposure (e–/Å^2^) | 2.3 | 2.3 |  |  |
| Defocus range (μm) | -1.0 to -3.0 | -1.0 to -3.0 |  |  |
| Pixel size (Å) | 3.17 | 1.57 |  |  |
| Images  acquired (no.) | 10067 | 7011 |  |  |
| Acquisition software | TFS EPU 2 | TFS EPU 2 |  |  |
| Symmetry imposed | I | I |  |  |
| Initial particle images (no.) | 230,000 | 2,725,140 | 750,242 | 17,549 |
| Final particle images (no.) | 65,000 | 1,020420 | 627,748 | 17,549 |
| Map resolution (Å)  FSC threshold | 6.4  0.143 | 3.77  0.143 | 6.62  0.143 | 18.13  0.143 |
| Map B-factor | 188.3 | 198.2 | 559.3 | 360.2 |
| **Refinement** |  |  |  |  |
| Initial model used (PDB code) | - | 1m1c | - | - |
| Map sharpening *B* factor (Å^2^) | - | 0 (not modified) | - | - |
| Model composition  Non-hydrogen atoms  Protein residues | -  - | 10302  1302 |  |  |
| *B* factors  Protein (min/max/mean) | - | 76.63/187.82/113.84 |  |  |
| R.m.s. deviations  Bond lengths (Å)  Bond angles (°) | -  - | 0.004 (0)  0.985 (0) |  |  |
| Validation  MolProbity score  Clashscore  Poor rotamers (%) | -  -  - | 2.12  15.89  0.45 |  |  |
| Ramachandran plot  Favored (%)  Allowed (%)  Disallowed (%) | -  -  - | 93.68  6.32  0.00 |  |  |
| Map-CC | - | 0.84 |  |  |

*Part of the dataset was acquired at 13 frames; the last frame was removed during image analysis.

**Supplementary Table 2. Identification of structural homologues of L-A virus region 481-587:** Structural alignment was performed by DALI server (http://ekhidna2.biocenter.helsinki.fi/dali/). Z score (Z), root-mean-square-deviation (RMSD) between the two aligned structures, the length of the aligned residues (lali) and the sequence identity (%id) are displayed. The description is fetched from the PDB entry.

| Chain | Z | rmsd | lali | %id | Description |
| --- | --- | --- | --- | --- | --- |
| 1m1c-B | 14.9 | 1.1 | 97 | 100 | Major Coat Protein |
| 1m1c-A | 7.6 | 0.8 | 97 | 100 | Major Coat Protein |
| 3e4g-A | 3.5 | 3.7 | 64 | 9 | ATP Synthase Subunit S, Mitochondrial |
| 3e2j-A | 3.5 | 3.8 | 61 | 10 | ATP Synthase Subunit S, Mitochondrial |
| 6kms-C | 3.4 | 3.2 | 46 | 9 | Methyltransferase N6AMT1 |
| 3dze-A | 3.3 | 3.7 | 65 | 9 | ATP Synthase Subunit S, Mitochondrial |
| 6k0x-A | 3.3 | 3.5 | 45 | 9 | Methyltransferase N6AMT1 |
| 2dfx-I | 3.2 | 2.8 | 60 | 7 | Colicin-E5 |
| 2fhz-A | 3.2 | 2.8 | 60 | 7 | Colicin-E5 Immunity Protein |
| 3qph-A | 3.1 | 3.3 | 61 | 7 | TRMB, A Global Transcription Regulator |
| 3wvq-A | 3.1 | 3.8 | 65 | 17 | PGM1 |
| 3wvr-B | 3.1 | 3.8 | 65 | 17 | PGM1 |
| 6khs-A | 3.1 | 3.1 | 45 | 11 | Methyltransferase N6AMT1 |
